# The deubiquitinase USP9X regulates RIT1 protein abundance and oncogenic phenotypes

**DOI:** 10.1101/2023.11.30.569313

**Authors:** Amanda K. Riley, Michael Grant, Aidan Snell, Athea Vichas, Sitapriya Moorthi, Anatoly Urisman, Pau Castel, Lixin Wan, Alice H. Berger

## Abstract

*RIT1* is a rare and understudied oncogene in lung cancer. Despite structural similarity to other RAS GTPase proteins such as KRAS, oncogenic RIT1 activity does not appear to be tightly regulated by nucleotide exchange or hydrolysis. Instead, there is a growing understanding that the protein abundance of RIT1 is important for its regulation and function. We previously identified the deubiquitinase *USP9X* as a RIT1 dependency in *RIT1*-mutant cells. Here, we demonstrate that both wild-type and mutant forms of RIT1 are substrates of USP9X. Depletion of USP9X leads to decreased RIT1 protein stability and abundance and resensitizes cells to EGFR tyrosine kinase inhibitors. Our work expands upon the current understanding of RIT1 protein regulation and presents USP9X as a key regulator of RIT1-driven oncogenic phenotypes.

## INTRODUCTION

Lung cancer remains the leading cause of cancer-related deaths in the United States and globally^1^. Non-small cell lung cancer (NSCLC) is the most prevalent type of lung cancer, and lung adenocarcinoma (LUAD) is the most common histological subtype of NSCLC^2^. Given that approximately 238,000 individuals will be diagnosed with lung cancer in 2023^1^, even rare molecular subtypes, defined by biomarkers found in ≤5% of tumors^3^, affect tens of thousands of individuals every year. In 2014, *RIT1* (Ras-like in all tissues) was identified as a rare LUAD oncogene that activates RAS signaling in tumors without canonical *EGFR* or *KRAS* mutations^4,5^. *RIT1* is mutated in 2% of LUAD tumors and amplified in another 14%^4,5^. The biological effect of *RIT1* amplifications in lung cancer is not well understood, but recent work suggests that *RIT1* amplifications phenocopy mutant *RIT1*^6^. In addition to LUAD, *RIT1* alterations have been identified in other cancers such as myeloid malignancies, uterine carcinosarcoma, and hepatocellular carcinoma^7–9^. Additionally, germline variants in *RIT1* cause Noonan syndrome^10^, an inherited “RAS-opathy” characterized by cardiac abnormalities and differences in craniofacial morphology.

*RIT1* is a RAS-related small GTPase with structural similarity to KRAS^4,11,12^. Similarly to other RAS proteins, RIT1 cycles between GDP- and GTP-bound states and can activate downstream MAPK (Mitogen-activated protein kinase) signaling when bound to GTP^4,13,14^. Although RIT1 is known to affect MAPK signaling, this regulation differs across cell types^4,14,15^. Of the currently known mutations, the M90I variant (RIT1^M90I^) is most prevalent in lung cancer, although there is a diversity of other somatic variants that occur, mostly clustering near M90I in the switch II domain^4^. *RIT1* mutants have similar or increased GTP loading compared to wild-type tightly regulated by GAPs and GEFs; mutations in *KRAS* impair this regulation and substantially increase the GTP-to-GDP ratio of KRAS^16^. *RIT1* mutations, however, appear to ablate regulation of RIT1 protein abundance^14^. Wild-type RIT1 can be polyubiquitinated and targeted for proteasomal degradation by the CUL3 RING E3 ligase and the adaptor protein LZTR1 (leucine zipper-like transcription regulator 1), but mutant forms of RIT1 evade this regulation^14^. LZTR1 appears to be a critical regulator of RIT1 in Noonan Syndrome because homozygous inactivating mutations in LZTR1 lead to increased RIT1 expression and, like mutant RIT1, cause Noonan syndrome^14,17,18^. This mechanism is not exclusive to Noonan Syndrome and has also been explored in the context of cancer. *LZTR1* mutations are found in myeloid malignancies, and *LZTR1* knockout increases the abundance of RIT1 in these cells , thereby promoting increased phosphorylation of MEK1/2 and ERK1/2^7,19,20^.

Recent work suggests that mass action of mutant RIT1 molecules recruits RAF kinases to the plasma membrane and activates RAS-related signaling and cell proliferation^21^. Increased abundance of RIT1^M90I^ is mediated by resistance to LZTR1-mediated degradation^14^, but it is not yet clear how over-expression of wild-type RIT1 is maintained within the cell. Multiomic analysis that examined proteomic, phosphoproteomic, and transcriptomic datasets revealed that over-expression of wild-type RIT1 phenocopies RIT1^M90I^ in terms of gene expression and activation of effector pathways^6^. Our current understanding of RIT1 oncogenesis is converging on the notion that protein abundance is crucial for cancer progression driven by RIT1; however, in lung adenocarcinoma it is not clear how RIT1 protein abundance or other mechanisms contribute to tumor growth.

In order to better understand RIT1 biology and genetic dependencies, we previously designed a CRISPR screening approach to identify genes required for RIT1^M90I^-driven resistance to EGFR inhibitors such as erlotinib and osimertinib^15,22^. To date, chemotherapy is the only treatment option for *RIT1*-driven cancers; therefore, identification of additional genetic vulnerabilities could be a key step in developing new targeted therapies and personalized medicine approaches. Furthermore, resistance to EGFR inhibitors remains a major clinical problem. Upwards of 35% of patients are predicted to harbor intrinsic resistance mutations, and acquired resistance often develops 1-2 years post-treatment^23,24^. Therefore, our CRISPR screening strategy also provides important insight to better understand the biology of TKI resistance.

To identify additional regulators of RIT1 abundance beyond LZTR1, we mined RIT1 dependencies for ubiquitin pathway-related genes and identified the deubiquitinase (DUB) USP9X as a candidate. As a DUB, USP9X removes polyubiquitin chains and prevents the degradation of protein substrates^25–30^. Here, we demonstrate that both wild-type and mutant RIT1 are substrates of USP9X. This work expands upon our understanding of RIT1 biology and presents USP9X as a potentially important clinical target for future studies.

## RESULTS

### *USP9X* is an essential gene in *RIT1*-mutant cells

To identify potential regulators of RIT1 protein abundance, we analyzed data from our previous CRISPR screen of genetic dependencies of RIT1^M90I^. In the prior work, we identified genes that were required for RIT1^M90I^ to promote resistance to the EGFR tyrosine kinase inhibitor (TKI) erlotinib in *EGFR*-mutant PC9 lung adenocarcinoma cells^15^ (**Figure 1A**). When comparing the CRISPR scores in erlotinib-treated vs. DMSO-treated screens, *RIT1* emerged as a top essential gene, as expected (**Figures 1B, S1A, and Supplementary Table 1**). Another top essential gene was the deubiquitinase *USP9X* (**Figures 1B, S1A, and Supplementary Table 1**). USP9X removes polyubiquitin chains to stabilize protein substrates and prevent their proteasomal degradation^25–30^. In cancer, *USP9X* has been identified as a tumor suppressor and an oncogene, depending on the cellular context^25,31–33^. To validate the result of the screen that USP9X is necessary for RIT1-induced resistance to EGFR inhibition, we generated pooled populations of RIT1^M90I^-mutant PC9-Cas9 cells harboring a guide RNA targeting *USP9X* or *RIT1* (**Figure 1C**). Knockout of *USP9X* or *RIT1* resensitized cells to erlotinib and osimertinib (**Figures 1D-E and S1B-C**). As an orthogonal approach to CRISPR knockout, we utilized siRNAs to knockdown *USP9X*. Knockdown of *USP9X* resensitized RIT1^M90I^-mutant cells to erlotinib (**Figure 1F**) and osimertinib (**Figure S1D**). Together, these experiments show that USP9X is required for *RIT1*-driven drug resistance in RIT1^M90I^-mutant PC9 cells.

**Figure 1.**
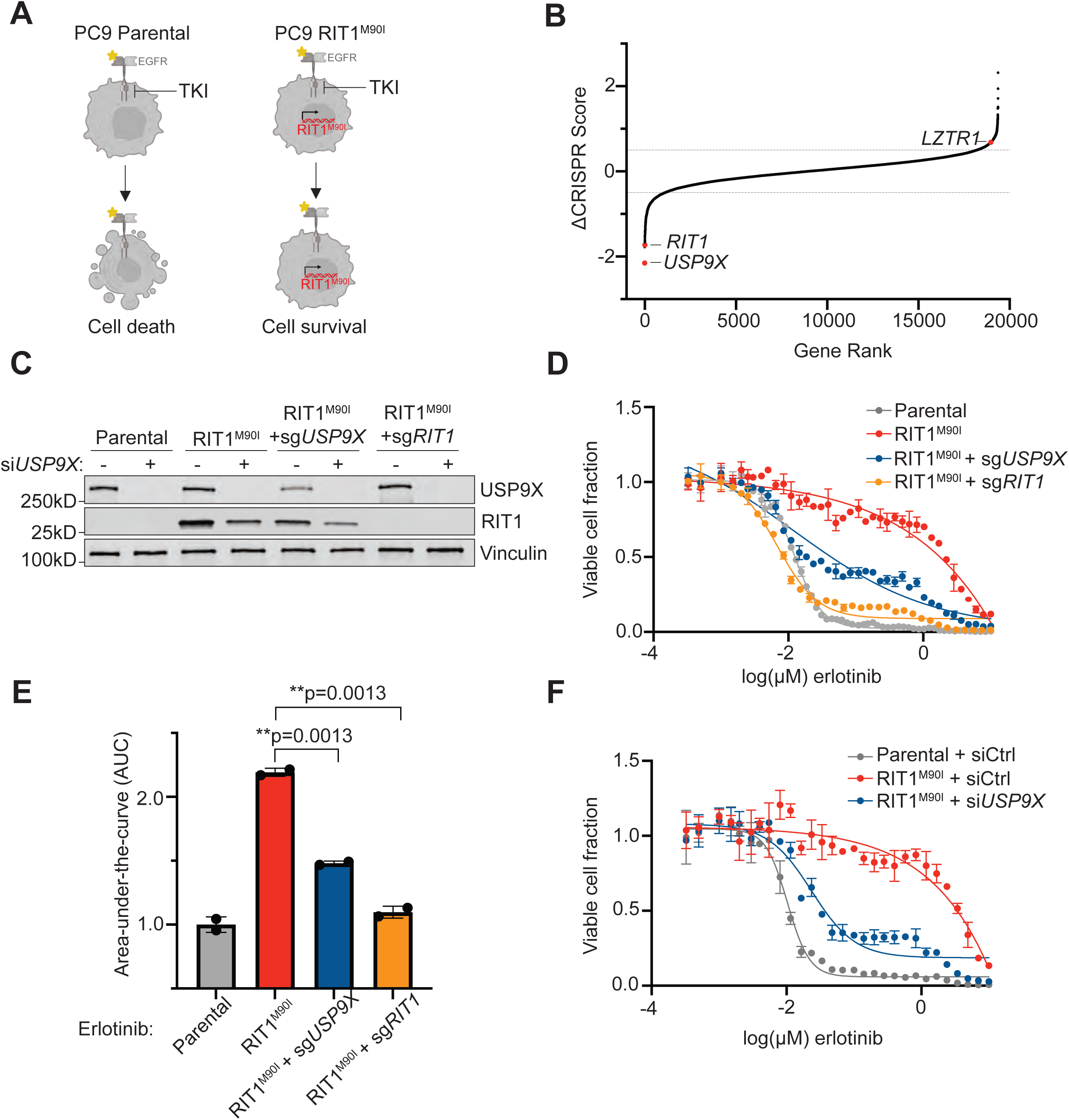
USP9X depletion reverses *RIT1*-driven drug resistance. **A,** Schematic of *RIT1*-driven EGFR tyrosine kinase inhibitor (TKI) resistance. Left, *EGFR*-mutant PC9 cells are sensitive to EGFR TKI’s such as erlotinib. Right, expression of RIT1^M90I^ in PC9 cells confers EGFR TKI resistance. Figure created with Biorender.com. **B,** Gene rank plot of previously published CRISPR/Cas9 whole-genome screen performed in RIT1^M90I^-mutant PC9-Cas9 cells. ΔCRISPR Score is the difference between CRISPR scores in the erlotinib screen vs. CRISPR Scores in the DMSO screen. **C,** Western blot of PC9-Cas9 cells treated with siCtrl or si*USP9X* for 48 hours. Vinculin serves as a loading control. **D,** Dose-response curves of PC9-Cas9 Parental cells and RIT1^M90I^-mutant PC9-Cas9 cells with indicated gene knockouts (sg*RIT1* and sg*USP9X*) treated with erlotinib for 72 hours. Knockouts confirmed from Western blot in (C). CellTiterGlo was used to quantify viable cell fraction determined by normalization to DMSO control. Data shown are the mean ± s.d. of two technical replicates. Data are representative results from n = 2 independent experiments. **E,** Area-under-the-curve (AUC) analysis of dose response curves shown in (D). p-values calculated by unpaired two-tailed t-tests. **F,** Dose-response curves of RIT1^M90I^-mutant PC9 cells treated with siCtrl or si*USP9X* for 48 hours, prior to treatment with erlotinib for 72 hours. Knockdowns validated by Western blot in (C). CellTiterGlo was used to quantify viable cell fraction determined by normalization to DMSO control. Data shown are the mean ± s.d. of two technical replicates. Data are representative results from n = 3 independent experiments.

Given that the protein abundance of RIT1 is known to be important for its function^14^, we were intrigued to see the deubiquitinase *USP9X* as a top essential gene in *RIT1*-mutant cells (**Figures 1B, S1A, and Supplementary Table 1**). We hypothesized that USP9X may be positively regulating RIT1 levels, and *USP9X* knockout would reduce RIT1 protein abundance. Indeed, *USP9X* knockout reduced RIT1^M90I^ protein abundance, and complete ablation of *USP9X* with an additional siRNA resulted in further reduction of RIT1^M90I^ (**Figure 1C**). This suggests that USP9X positively regulates RIT1 protein abundance.

In addition to EGFR TKI resistance, expression of RIT1^M90I^ is known to promote anchorage-independent growth^4^. Given this, we investigated how USP9X regulates proliferation phenotypes in *RIT1*-mutant PC9 cells. Under normal media conditions, genetic depletion of *RIT1* and *USP9X* did not affect the proliferation of *RIT1*-mutant cells (**Figure S2**). In the context of erlotinib, PC9-Cas9-RIT1^M90I^ cells depend on RIT1 for growth; therefore, genes required under erlotinib treatment are RIT1 dependency genes (**Figure 1A**). Because of this, we expect that the effect of *USP9X* depletion will be most pronounced when cells are treated with an EGFR inhibitor. Under erlotinib treatment, RIT1^M90I^ + sg*USP9X* cells and RIT1^M90I^ + sg*RIT1* cells proliferated slower than RIT1^M90I^ cells (**Figure 2A**). In addition to 2D growth, we explored 3D growth via soft agar colony formation assays in DMSO and erlotinib (**Figure 2B**). As expected, in DMSO (vehicle), there was no difference in the number of colonies formed by RIT1^M90I^ + sg*USP9X* cells or RIT1^M90I^ + sg*RIT1* cells compared to parental or RIT1^M90I^-mutant cells (**Figure 2C**), although RIT1 appeared to influence colony size in these conditions (**Figure 2D**). However, in the presence of EGFR inhibition with erlotinib, RIT1^M90I^ induced 738-fold more colony formation than parental cells, and this colony formation was almost completely ablated by *RIT1* or *USP9X* knockout (**Figure 2E**). Of the colonies that formed in this condition, all were significantly smaller than those formed by RIT1^M90I^-mutant cells (**Figure 2F**). These data demonstrate that USP9X is important for maintaining *RIT1*-driven proliferation and anchorage-independent growth.

**Figure 2.**
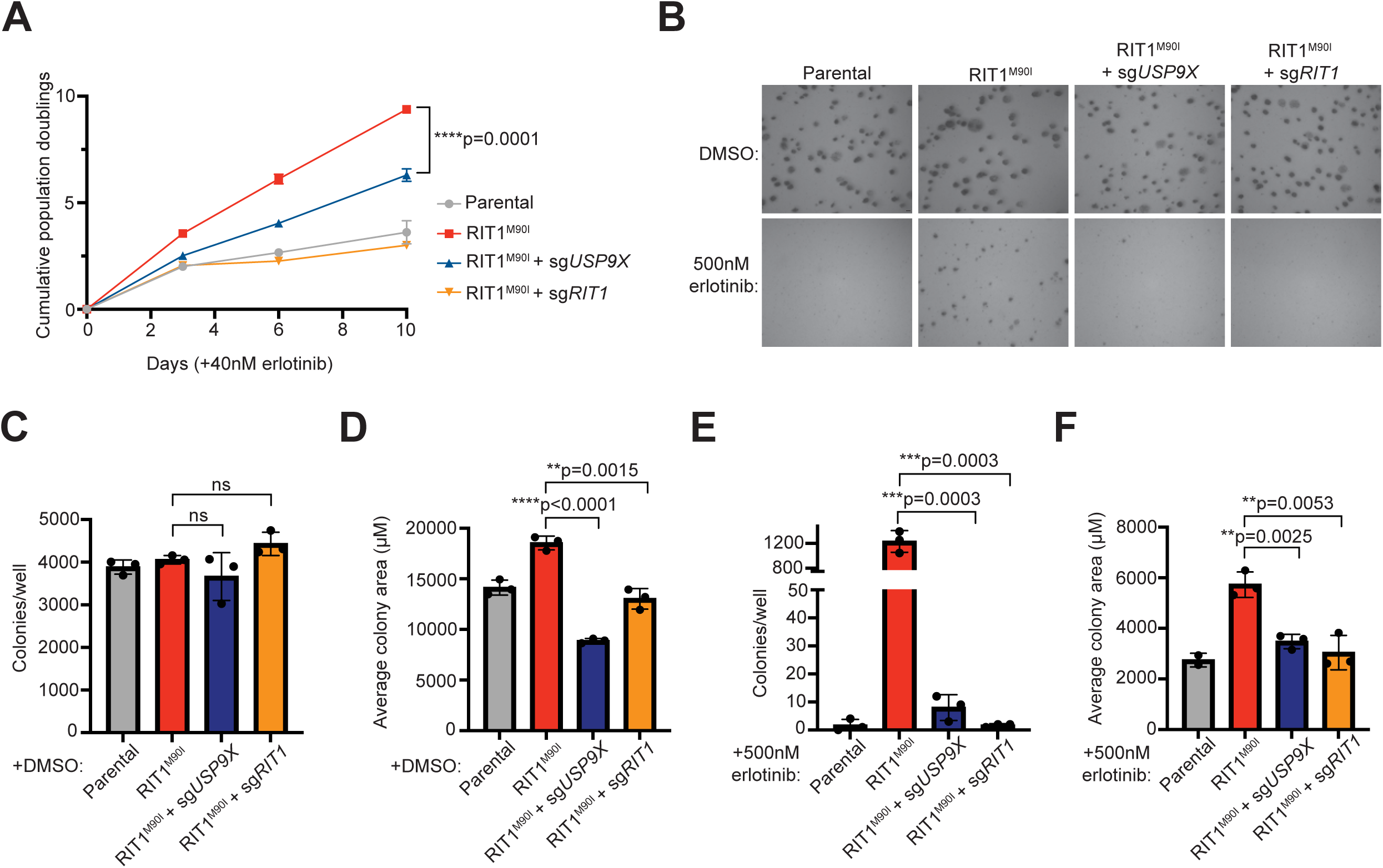
USP9X regulates *RIT1*-driven proliferation and anchorage-independent growth. **A,** Proliferation of PC9-Cas9 Parental cells and RIT1^M90I^-mutant PC9-Cas9 cells with indicated gene knockouts (sg*RIT1* and sg*USP9X*) treated with 40 nM erlotinib. Data shown are the mean ± s.d. of three technical replicates per cell line. Data are representative results from n = 2 independent experiments. p-value calculated by multiple unpaired two-tailed t-tests. **B,** Representative images of soft agar colony formation assay in PC9-Cas9 (Parental) cells and RIT1^M90I^-mutant PC9-Cas9 cells with indicated gene knockouts (sg*RIT1* and sg*USP9X*) treated with DMSO or 500 nM erlotinib. Images captured at 4X after 10 days of growth. Scale bar is 100 μM. **C,** Counts of colonies per well formed by indicated cell lines treated with DMSO. Counts taken after 10 days of growth. Data shown are the mean ± s.d. of three technical replicates per cell line. ns = not significant by unpaired two-tailed t-test. **D,** Average colony area of all colonies formed by indicated cell lines treated with DMSO for 10 days. Data shown are the mean ± s.d. of three technical replicates per cell line. p-values calculated by unpaired two-tailed t-tests. **E,** Counts of colonies per well formed by indicated cell lines treated with 500 nM erlotinib for 10 days. Data shown are the mean ± s.d. of three technical replicates per cell line. p-values calculated by unpaired two-tailed t-tests. **F,** Average colony area of all colonies formed by indicated cell lines treated with 500 nM erlotinib for 10 days. Data shown are the mean ± s.d. of three technical replicates per cell line. p-values calculated by unpaired two-tailed t-tests. Data in C-F are representative results from n = 2 independent experiments.

### USP9X regulates RIT1 abundance and stability in multiple cell lines

We hypothesized that *USP9X* knockout may reduce the abundance of RIT1, thus explaining USP9X’s ability to counteract RIT1 function observed above. We already observed that *USP9X* knockout reduced RIT1 abundance in RIT1^M90I^-mutant PC9 cells (**Figure 1C**), and we expanded upon these findings. In parental PC9 cells (which express endogenous, wild-type RIT1), si*USP9X* significantly reduced wild-type RIT1 protein abundance (**Figure 3A-B**). In PC9 cells expressing ectopic RIT1^M90I^, CRISPR-mediated depletion of *USP9X* also significantly reduced the abundance of RIT1^M90I^ (**Figure 3C-D**). In addition to protein abundance, we explored the stability of RIT1 in the context of *USP9X* depletion. Cells were treated with cycloheximide (CHX) to inhibit protein translation, and the level of RIT1 was monitored over time by western blot. In *USP9X* knockout PC9 cells, RIT1 level decreased faster over the course of 12 hours compared to parental cells (**Figure 3E**). The half-life of RIT1 in parental cells was approximately 7.6 hours compared to 2.4 hours in *USP9X* knockout cells (**Figure 3F**). By the 6 hour time-point, we consistently found that RIT1 protein abundance was significantly lower in *USP9X* knockout cells compared to parental (**Figure 3G**). Together, these data show that USP9X is important for maintaining RIT1 protein abundance.

**Figure 3.**
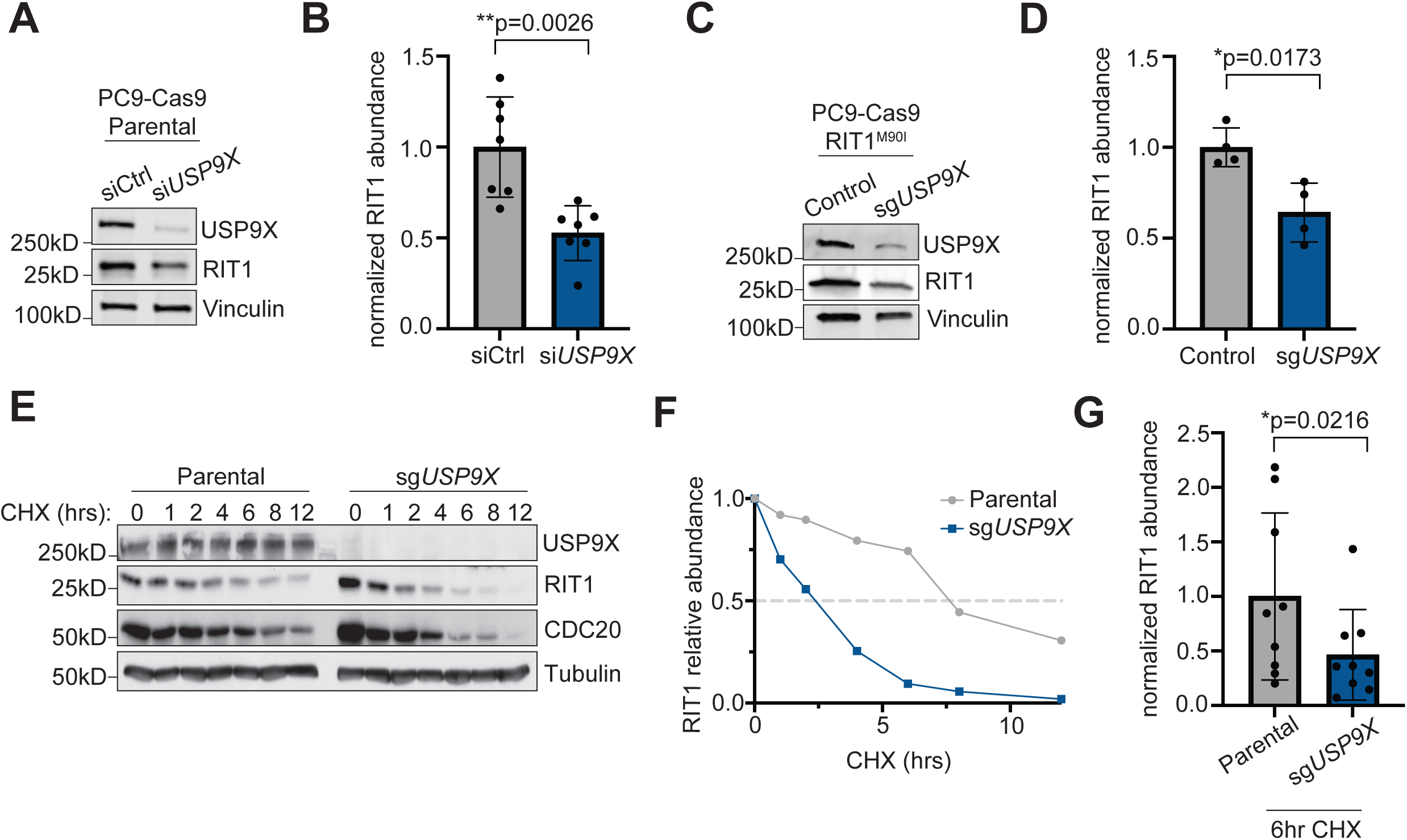
USP9X controls RIT1 abundance and stability in PC9 lung adenocarcinoma cells. **A,** Western blot of PC9-Cas9 Parental cells treated with indicated siRNAs for 48 hours. Vinculin serves as a loading control. **B,** Quantification of Western blot band intensity of RIT1 bands in (A). Data shown are the mean ± s.d. of three independent experiments with 2-3 biological replicates per condition. p-value calculated by paired two-tailed t-test. **C,** Western blot of RIT1^M90I^-mutant PC9-Cas9 cells (Control) and a CRISPR-engineered clonal *USP9X* KO (sg*USP9X*) cell line. Vinculin serves as a loading control. **D,** Quantification of Western blot band intensity of RIT1 bands in (C). Data shown are the mean ± s.d. of two independent experiments with 2 biological replicates per condition. p-value calculated by paired two-tailed t-test. **E,** PC9-Cas9 Parental and sg*USP9X* cells treated with 100 μg/mL cycloheximide (CHX) for the indicated time periods before harvesting for Western blot. CDC20 serves as a positive control for USP9X activity. Tubulin serves as a loading control. Data are representative of n = 3 independent experiments. **F,** Half-life analysis of RIT1 protein abundance over time based on RIT1 band intensity in (E). **G,** Comparison of RIT1 protein abundance based on Western blot band intensity of aggregated cycloheximide-chase experiments in cells treated with CHX for 6 hours. Data shown are the mean ± s.d. of three independent experiments with 3 technical replicates per cell line. p-value calculated by paired two-tailed t-test.

To extend these findings to another cell context, we utilized NCI-H2110 cells, which is currently the only commercially available lung adenocarcinoma cell line that harbors an endogenous RIT1^M90I^ mutation^4^. Similar to our results in PC9 cells, treatment with si*USP9X* significantly reduced the abundance of RIT1^M90I^ (**Figure S3A-B**). As orthogonal confirmation, we generated NCI-H2110 cell lines incorporating an inducible (iCas9) system where expression of Cas9 is under control of doxycycline (dox)^34^. Following dox treatment and Cas9 expression, *USP9X* knockout significantly decreased RIT1 abundance (**Figure S3C-D**). To confirm that USP9X-mediated regulation of RIT1 is occurring at the protein level, we performed RT-qPCR. No difference in *RIT1* mRNA expression was observed in sg*USP9X* cells (**Figure S3E**), indicating that USP9X’s regulation of RIT1 occurs post-transcriptionally and occurs in multiple cell contexts.

### RIT1 ubiquitination is mediated by USP9X’s catalytic activity

To explore USP9X’s regulation of RIT1 in an unbiased fashion and to identify other potential DUBs of RIT1, we fused RIT1 to a ubiquitin molecule through a flexible peptide linker to obtain a constitutively ubiquitinated form of RIT1 that we termed RIT1^∼Ub^ (**Figure 4A**). This construct acts as a molecular trap, by stabilizing interaction with DUBs, which are unable to cleave the peptide bond^35^. Next, we undertook affinity-purification mass spectrometry in HEK293T cells using our RIT1^∼Ub^ mutant. In cells expressing RIT1^∼Ub^, we found enrichment of RIT1 and LZTR1 peptides compared to control cells transfected with empty vector (**Figures 4B and S4A-C**). We also found a similar enrichment of USP9X peptides (**Figures 4B, S4A and S4D**), indicating that USP9X is physically interacting with ubiquitinated RIT1.

**Figure 4.**
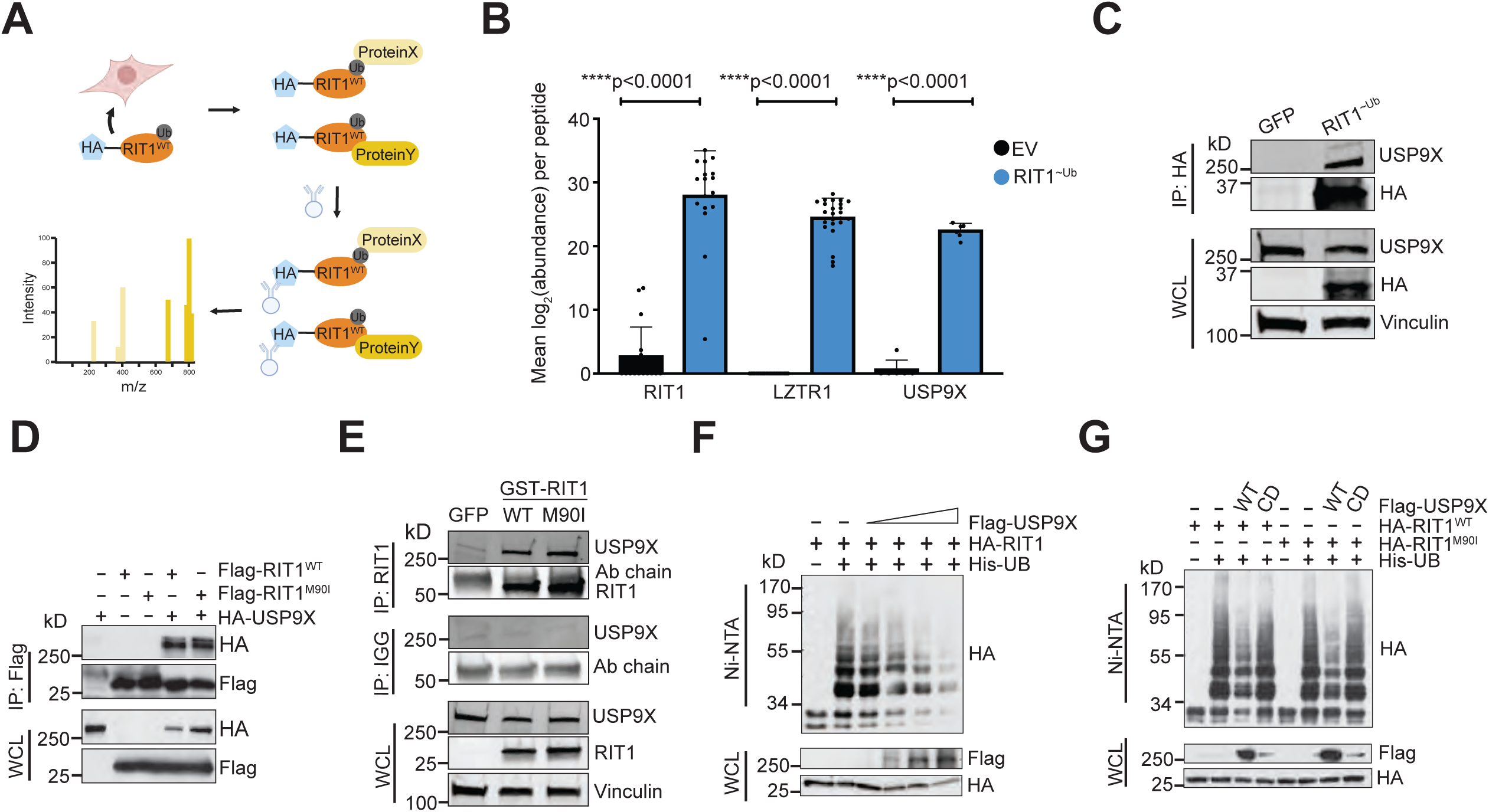
USP9X binds to and deubiquitinates RIT1. **A,** Schematic of affinity-purification/mass-spectrometry (AP/MS) experiment performed in HEK293T cells transfected with RIT1^∼Ub^ vector. This experiment was designed to identify proteins that bind to ubiquitinated RIT1. Figure created with Biorender.com. **B,** Mean abundance (log_2_-transformed) of peptides across biological replicates (n = 7 for Empty Vector/EV and n = 4 for RIT1^∼Ub^) of affinity purification/mass spectrometry experiment. p-values calculated by paired two-tailed t-tests. **C,** Co-immunoprecipitation in HEK293T cells transfected with GFP control or RIT1^∼Ub^. Vinculin serves as a loading control. Data shown is representative of n = 2 independent experiments. **D,** Western blot of whole cell lysates (WCL) and anti-Flag immunoprecipitates (IP) derived from HEK293T cells transfected with Flag-RIT1^WT^ or Flag-RIT1^M90I^ together with the HA-USP9X construct. 36 hours post-transfection, cells were pretreated with 10 μM MG132 for 10 hours before harvesting. Data shown are representative of n = 4 replicates for RIT1^WT^ and n = 1 replicates for RIT1^M90I^. **E,** Co-immunoprecipitation experiment in HEK293T cells transfected with indicated GST-tagged RIT1 variants or a GFP transfection control. Vinculin serves as a loading control. Data shown are representative of n = 2 independent experiments. **F,** Western Blot of WCL and subsequent His-tag pull-down in 6 M guanine-HCl containing buffer derived from HEK293T cells transfected with the indicated plasmids. Cells were pretreated with 10 μM MG132 for 16 hours to block the proteasome pathway before harvesting. Data shown are representative of n = 3 independent experiments. **G**, Ubiquitination experiment as described in (F) in HEK293T cells transfected with RIT1^WT^ and RIT1^M90I^, as well as wild-type or catalytically dead (CD) USP9X. Data shown are representative of n = 3 independent experiments.

As an orthogonal approach to our AP/MS method to assess the physical interaction of USP9X and RIT1, we performed co-immunoprecipitation experiments in HEK293T cells expressing FLAG-tagged RIT1 (wild-type and RIT1^M90I^) and HA-tagged USP9X. As expected, we saw evidence for interaction of RIT1^WT^ and RIT1^M90I^ with USP9X (**Figure 4D**). To rule out potential confounding factors related to tagged USP9X, we performed this experiment with endogenous USP9X in HEK293T cells and found that endogenous USP9X also interacts with RIT1^WT^ and RIT1^M90I^ (**Figure 4E**).

To investigate if USP9X is regulating RIT1 ubiquitination, we transiently transfected HA-tagged RIT1 and His-tagged ubiquitin in HEK293T cells. We titrated increasing amounts of Flag-USP9X and found that ubiquitinated RIT1 decreased in a dose-dependent manner in relation to the amount of USP9X expressed (**Figure 4F**). To expand upon these findings, we performed ubiquitin-pulldown experiments with a catalytically dead (CD) form of USP9X that harbors an active site mutation (C1566A) which ablates its deubiquitinase activity^30^. Expression of wild-type USP9X reduced ubiquitination of RIT1^WT^ and RIT1^M90I^, but expression of USP9X^CD^ did not affect RIT1 ubiquitination (**Figure 4G**). These experiments confirm that the deubiquitinase activity of USP9X is responsible for modulating ubiquitination of RIT1.

### USP9X could be a promising therapeutic target for *RIT1*-driven diseases

Our findings suggest that *USP9X* genetic depletion reduces RIT1 abundance and abrogates *RIT1*-driven oncogenic phenotypes. As such, pharmacological inhibition of USP9X is predicted to have similar effects, and USP9X could be a promising drug target in diseases characterized by *RIT1* mutations and amplifications. Of note, these implications are not limited to LUAD. Analysis of the Cancer Dependency Map and associated proteomics datasets^36^ revealed a positive correlation between USP9X protein abundance and RIT1 protein abundance (**Figure 5A**), suggesting that this regulation may extend to multiple cancer types driven by *RIT1* alterations. Importantly, this correlation was also seen with CDC20–a known USP9X substrate^30^ (**Figure 5B**), whereas no correlation was observed when comparing USP9X protein abundance to mRNA expression of *RIT1* (**Figure 5C**) or *CDC20* (**Figure 5D**)^37^. Thus, we propose pharmacological inhibition of USP9X as a strategy to promote RIT1 degradation, which would be detrimental to the growth and proliferation of *RIT1*-driven tumors (**Figure 5E**).

**Figure 5.**
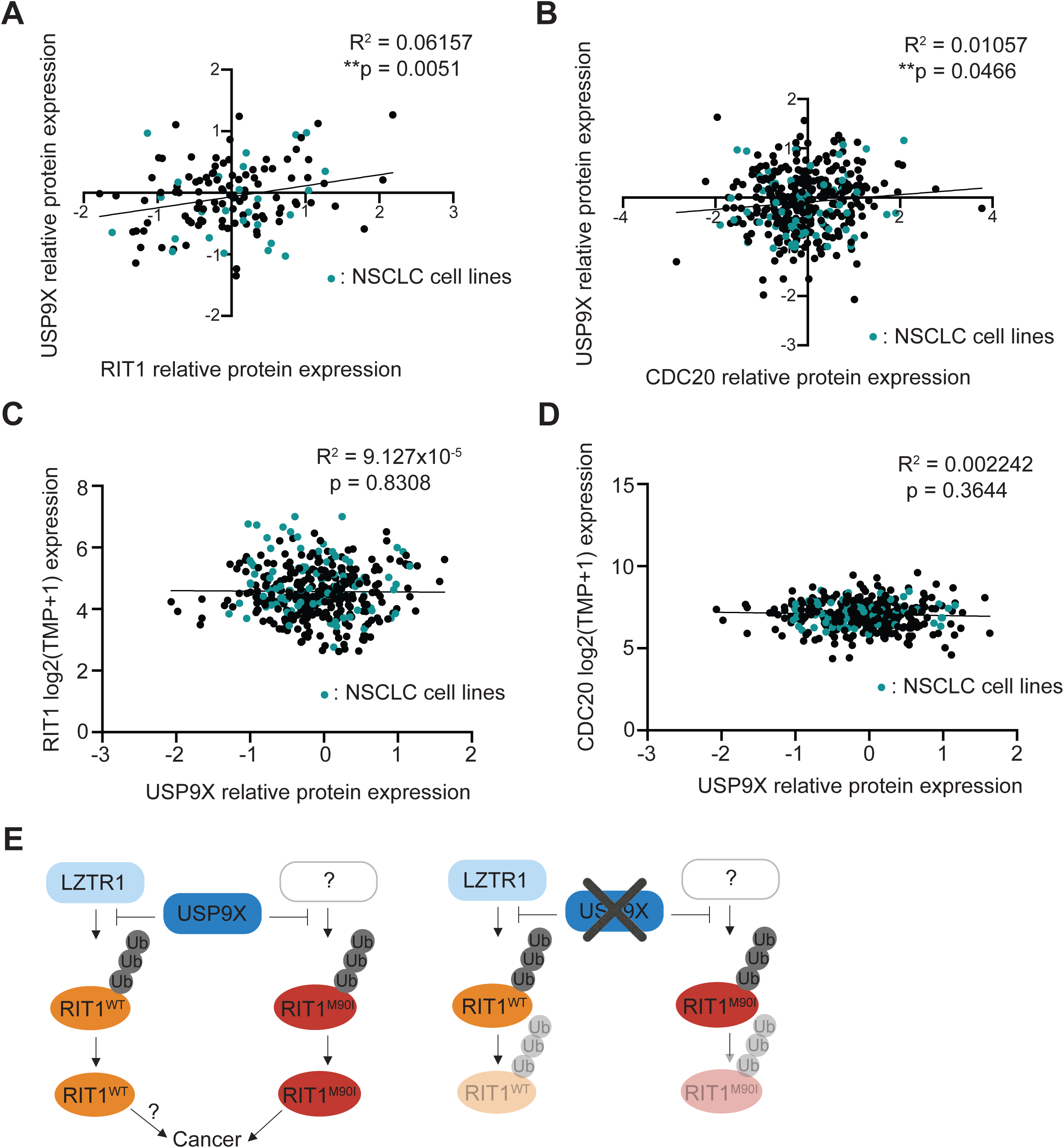
USP9X-mediated regulation of RIT1 is relevant across cancer types. **A,** Correlation of proteomics data^36^ from the Cancer Dependency Map (DepMap) comparing USP9X (Q93008) and RIT1 (Q92963-3). Pearson R squared and p-values calculated in Prism. **B,** Correlation of DepMap proteomics data^36^ comparing USP9X (Q93008) and CDC20 (Q12834). Pearson R squared and p-values calculated in Prism. **C,** Correlation of DepMap gene expression data (Expression Public 23Q2)^37^ for RIT1 and USP9X proteomics data (Q93008)^36^. Pearson R squared and p-values calculated in Prism. **D,** Correlation of DepMap gene expression data (Expression Public 23Q2)^37^ for CDC20 and USP9X proteomics data (Q93008)^36^. Pearson R squared and p-values calculated in Prism. **E,** Proposed model (left) of RIT1 protein regulation. RIT1^WT^ is ubiquitinated by LZTR1, while RIT1^M90I^ is ubiquitinated by a currently unknown E3 ligase. USP9X counteracts the ubiquitination of both wild-type and mutant RIT1. Increased RIT1 abundance and stability are important for RIT1 function and disease progression. The exact biological consequences of RIT1^WT^ amplification have yet to be elucidated. Genetic knockout (right) of *USP9X* prevents RIT1 deubiquitination, thereby promoting RIT1 degradation and abrogating oncogenic phenotypes. Figure created with Biorender.com.

Together, we propose a model whereby USP9X positively regulates wild-type and mutant RIT1 (**Figure 5E**). We predict that this regulation counteracts the effects of LZTR1 on wild-type RIT1 (**Figure 5E**). Mutant forms of RIT1 are known to evade regulation by the CRL3^LZTR1^ complex, which is composed of the CUL3 RING E3 ubiquitin ligase and the adaptor protein LZTR1^14^. The E3 ligase responsible for ubiquitinating mutant RIT1 has yet to be identified (**Figure 5E**). In summary, our work builds upon the understanding that the protein abundance of RIT1 is key to its function, and we have identified USP9X as a positive regulator of wild-type and mutant RIT1.

## DISCUSSION

Our group previously performed CRISPR screens to explore *RIT1* genetic dependencies^15^ and identified the deubiquitinase *USP9X* as a genetic regulator of RIT1 function^15^. This finding was particularly interesting given that there is a growing understanding that the protein abundance of RIT1 is important for its function^14,20,21^. Mutant forms of RIT1 evade regulation by the CRL3^LZTR1^ complex, thereby increasing RIT1 protein abundance^14^. As such, it is highly probable that the activity of RIT1 is also regulated by DUBs.

*USP9X*–first discovered in fruit flies (*Drosophila melanogaster*) as the *fat facets* gene–is essential for embryogenesis in flies and mice (*Mus musculus)*^38,39^. *USP9X* can replace *fat facets* during fly development^38^ and shares 44% identity and 88% similarity to the fruit fly gene^40^. Although USP9X’s catalytic domain is consistent with DUBs in yeast, *fat facets* is the earliest *USP9X* ortholog characterized^41^. From flies to mammals, *USP9X* is highly constrained across evolution^42^. Of note, orthologs of *RIT1* and *LZTR1* have been documented in fruit flies but not in less complex lab models such as *Caenorhabditis elegans*^43^. This contrasts with *KRAS*, which is highly conserved in yeast^43^. Given this, it is possible that a regulatory network involving *USP9X*, *RIT1*, and *LZTR1* has co-evolved from flies to mammals. Rigorous evolutionary analysis is required to support this hypothesis.

Although it was initially characterized in the context of development^44,45^, USP9X has been implicated in apoptosis^25,46,47^, protein trafficking^28,48–50^, and polarity^51–53^. USP9X has been found across a wide range of cellular compartments, including the cytoplasm^50^, nucleus^54,55^, and mitochondria^25^. This diversity highlights that USP9X function is largely dictated by cell type. As a deubiquitinase, USP9X positively maintains the abundance of proteins by removing polyubiquitin chains and preventing proteasomal degradation^25–30^. USP9X is known to remove polyubiquitin^25,56–58^ and monoubiquitin^28,59^ chains. Despite these diverse functions, structural analysis of USP9X suggests that the catalytic domain preferentially binds to and cleaves polyubiquitin chains with K48- and K11-linkages^56^. LZTR1 promotes K48 polyubiquitination of RIT1^WT^ at K187 and K135^14^. Given this, it is possible that USP9X deubiquitinates RIT1^WT^ at these lysine sites. Future work is needed to identify the precise lysines targeted by USP9X, and whether this varies between wild-type and mutant forms of RIT1.

In the context of cancer, *USP9X* can be upregulated or downregulated in various cancer types, depending on the target substrates^41^. In lung adenocarcinoma, *USP9X* has been found amplified^5,60^, deleted^5,60^, and mutated^60–64^. The exact consequences of these diverse mutations has yet to be fully elucidated. In NSCLC, *USP9X* has been characterized as an oncogene^65–67^, and high *USP9X* expression is associated with poorer overall survival^68^. Outside of cancer, *USP9X* mutations underlie X-linked developmental disability (XID)^69,70^. In the neurons of individuals afflicted with XID, *USP9X* knockout causes cytoskeletal disruptions that hinder cell growth and migration^69^. RIT1 is known to regulate neuronal growth and survival^71^ and can also affect actin dynamics in fibroblast-like cells via regulation of p21-activated kinase (PAK1)^72^. It is possible that USP9X-mediated regulation of RIT1 is important in the context of XID, but this requires further experimentation. These findings, in combination with the observation that USP9X can cleave a diverse range of ubiquitin linkages^56^, suggest that cellular context is important for determining key USP9X substrates and potential relevance in disease states.

Given the function of USP9X as a deubiquitinase, we predicted that USP9X would physically interact with and modify the ubiquitination status of RIT1. Our unbiased AP/MS approach revealed that USP9X interacts with mono-ubiquitinated RIT1 (**Figures 4A-B, S4A, and S4D**). As expected, this assay also detected LZTR1–a known RIT1 interactor–thereby increasing the robustness of our findings (**Figures 4B, S4A, and S4C**). We further validated the physical interaction of USP9X and RIT1 in HEK293T cells (**Figure 4D-E**) and confirmed that the catalytic activity of USP9X regulates the ubiquitination status of RIT1^WT^ and RIT1^M90I^ (**Figure 4F-G**). Our findings suggest that USP9X deubiquitinates RIT1^M90I^, but we have yet to identify the E3 ubiquitin ligase(s) that is promoting ubiquitination. In our CRISPR screen, we identified 22 E3 ligases that were positively selected (**Supplementary Table 1**), meaning that individual genetic knockout of these ligases conferred a growth advantage in *RIT1*-mutant PC9 cells. Therefore, these genes are candidate E3 ligases that may target RIT1^M90I^ independently or potentially cooperate with CRL3^LZTR1^ to ubiquitinate RIT1. Systematic characterization and investigation of these 22 E3 ligases is required to further elucidate the protein-level regulation of RIT1^M90I^.

Even though mutant RIT1 is the predominant form of RIT1 expressed in PC9 cells, LZTR1 was observed as a cooperating factor in our CRISPR screen^15^. In other words, knockout of *LZTR1* conferred a growth advantage in RIT1^M90I^-mutant PC9 cells (**Figures 1B, S1A, and Supplementary Table 1**). This result was somewhat unexpected given that the M90I mutation in RIT1 prevents its interaction with LZTR1^14^. However, it is possible that LZTR1 is acting on the endogenous wild-type RIT1 that is also expressed in our engineered *RIT1*-mutant PC9 cells. Our work predicts that both USP9X and LZTR1 are acting on wild-type RIT1, while only USP9X is targeting mutant RIT1 (**Figure 5E**). The findings presented here support currently known mechanisms of RIT1 regulation^14,20,43,73^, while also identifying USP9X as a novel regulator of both wild-type and mutant RIT1.

To confirm the dependency of *RIT1*-mutant cells on USP9X, we turned to our CRISPR screening system where we initially identified USP9X. This screen was performed with RIT1^M90I^-mutant PC9 cells, which depend on RIT1^M90I^ to confer resistance to EGFR inhibitors such as erlotinib or osimertinib (**Figure 1A**). In this setting, genes required under erlotinib treatment are RIT1 dependency genes. Given this context, we replicated the screen conditions when exploring USP9X-mediated regulation of RIT1. We validated this dependency in erlotinib- and osimertinib-treatment experiments (**Figures 1D-F and S1B-D**). Notably, complete knockdown of *USP9X* with siRNA resulted in a dramatic resensitization to erlotinib (**Figures 1C and 1F**) and osimertinib (**Figures 1C and S1D**), while incomplete knockout of *USP9X* via CRISPR editing only partially reverted this resistance phenotype (**Figures 1C-E and S1B-C**). These differences in sensitivity appear to be directly related to the knockout efficiency of various techniques and further support the conclusion that USP9X is important for regulating *RIT1*-driven drug resistance.

As expected, *USP9X* knockout impaired the proliferation of *RIT1*-mutant cells in erlotinib (**Figure 2A**). In soft agar colony formation assays, *USP9X* knockout cells formed fewer and smaller colonies (**Figure 2E-F**). These phenotypic effects appear to be directly related to USP9X’s regulation of RIT1 protein abundance. Overall, we consistently found that USP9X depletion corresponded with decreased RIT1 protein abundance (**Figures 1C**, **3A-D, and S3A-D**). Of note, we recognize the limitations of the PC9 cell system in studying RIT1. PC9 cells harbor a mutation in *EGFR*, but in patient tumors *RIT1* mutations are almost always mutually exclusive with other mutations in the RTK/RAS pathway^5^. Although the PC9+erlotinib/osimertinib system is an *in vitro*-based cell line model, it offers valuable insight into RIT1 regulation, genetic dependencies, and oncogenic mechanisms. Furthermore, our work in NCI-H2110 cells (**Figure S3**) and DepMap analyses (**Figure 5A-D**) underscores that RIT1 is a substrate of USP9X in other human lung cancer cell models and may be relevant across a wide range of cancer types. Indeed, USP9X could be a promising therapeutic target for diseases characterized by *RIT1* amplifications and mutations.

RIT1 regulation by a DUB could open opportunities to inhibit DUB function and thereby decrease RIT1 protein levels. Attempts have been made to develop small molecule inhibitors against USP9X. The compound WP1130 has been shown to inhibit USP9X as well as other DUBs including USP5, USP14, and UCH37^74^. In cells expressing high abundance of oncoproteins targeted by USP9X, WP1130 treatment abrogates growth and proliferation^31,75^. However, the pre-clinical practicality of WP1130 is limited due to low solubility and poor bioavailability in animal models^76,77^. The compound G9 is a newer USP9X inhibitor, and it is more soluble and less toxic than WP1130^77^. Experiments with G9 have shown promising results for breast cancer, leukemia, and melanoma cells harboring specific mutations^27,78–80^. Notably, G9 has also been shown to target USP24^81^ and USP5^82^, so it is difficult to directly link the effects of this drug to USP9X inhibition. In 2021, a more specific USP9X inhibitor FT709 was developed with a nanomolar range IC50^83^. Unlike WP1130 or G9, FT709 does not target USP24 or USP5^83^. It will be intriguing to test if FT709 destabilizes RIT1 in NSCLC cells and whether FT709 sensitizes EGFRi-resistant NSCLC cells to EGFR or MAPK inhibition.

In summary, we identified USP9X as a positive regulator of RIT1 function. USP9X deubiquitinates wild-type and mutant RIT1 (**Figure 4F-G**), thereby increasing RIT1 abundance and stability. Given that protein abundance of RIT1 is important for its function^14,20,21^, USP9X is a key factor in mediating *RIT1*-driven oncogenic phenotypes. Our work supports previously known mechanisms of RIT1 regulation by LZTR1^14,20,43^, and we suggest that USP9X and LZTR1 oppose the action of one another in controlling the ubiquitination status of wild-type RIT1 (**Figure 5E**). We found that USP9X also targets RIT1^M90I^ (**Figures 1C, 3C-D, 4D-E, 4G, and S3A-D**) and future work is needed to identify other players within the ubiquitin-proteasome system that may be regulating mutant forms of RIT1 (**Figure 5E and Supplementary Table 1**). Additionally, more experimentation is required to understand the biological consequences of RIT1^WT^ amplification in disease states. Overall, this work builds upon our knowledge of RIT1 biology and the mechanisms underlying how *RIT1* alterations cause disease and improves our understanding of the role of USP9X in lung cancer. These insights can be leveraged in the future to develop robust novel therapies for diseases characterized by *RIT1* mutations and amplifications.

## ACKNOWLEDGEMENTS

This research was funded in part through NCI F31CA271637 to A.K.R., NCI R37CA252050 and a Pardee Foundation award to A.H.B, R01CA255398 and ACS RSG-19-226-01-TBE to L.W., and ACS TLC-21-009-01-TLC to A.H.B. & L.W. This research was supported by the Cellular Imaging Shared Resource RRID:SCR_022609 of the Fred Hutch/University of Washington/Seattle Children’s Cancer Consortium (P30 CA015704). We would like to thank Dr. Lena Schroeder of the Fred Hutchinson Cancer Center Cellular Imaging Shared Resource for assistance with microscopy and image analysis. The wild-type and catalytically dead USP9X constructs used in ubiquitination experiments were a kind gift from Dr. Lindsey Allan at the University of Dundee and were previously published^30^.

## Author contributions

A.K.R., L.W., and A.H.B. conceived of and designed the study. A.K.R., M.G., and A.S. performed the cellular and biochemistry experiments. A.V. generated *RIT1*-mutant PC9-Cas9 knockout cell lines. S.M. analyzed the AP/MS data and generated associated plots. L.W. and A.S. provided panels 3E and 4D, and L.W. and M.G. provided panels 4F-G. P.C. contributed reagents. A.U. and P.C. performed AP/MS experiments and provided the associated methods. A.K.R. wrote the manuscript, which was reviewed by all co-authors.

## KEY RESOURCES TABLE

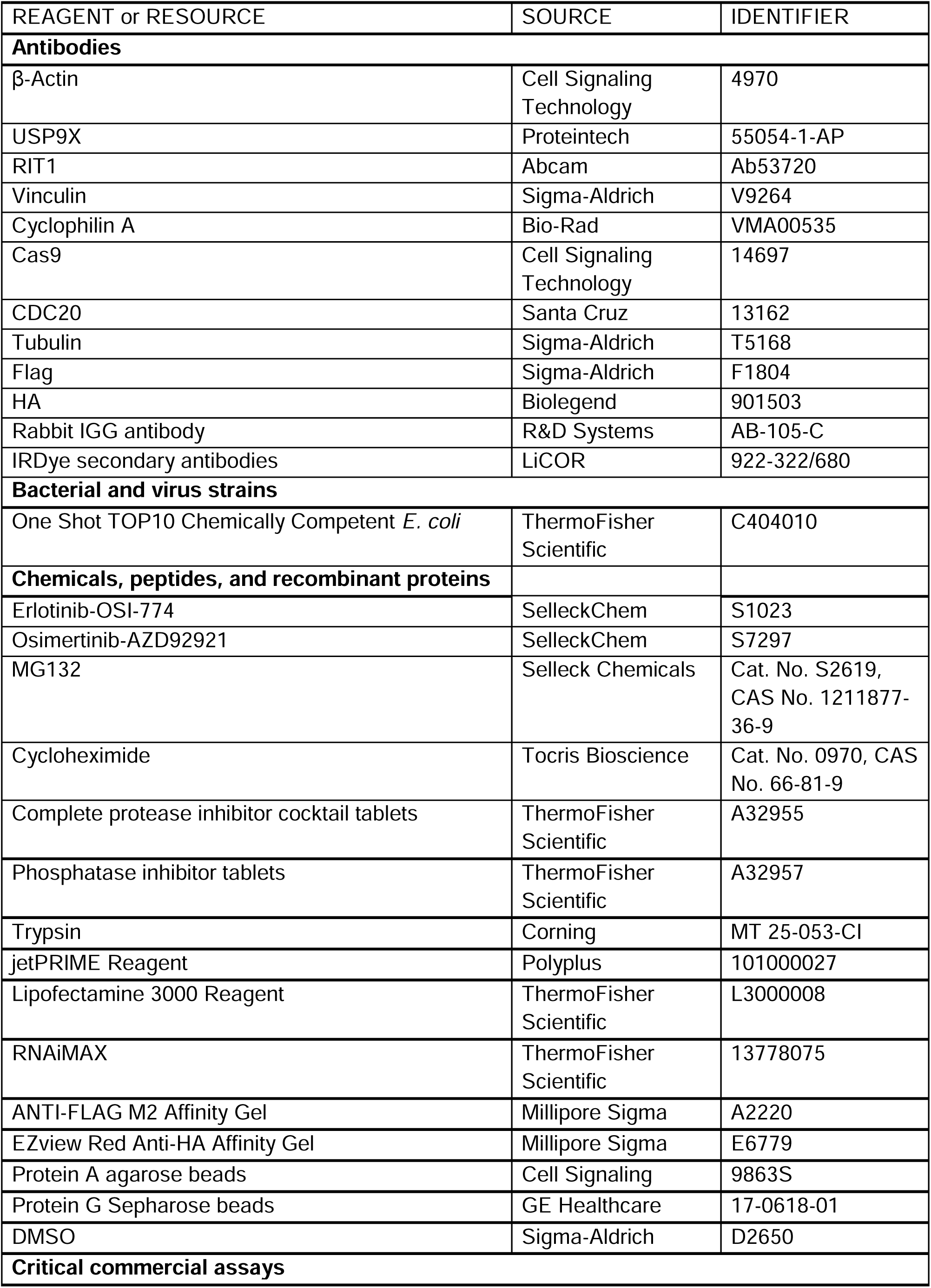
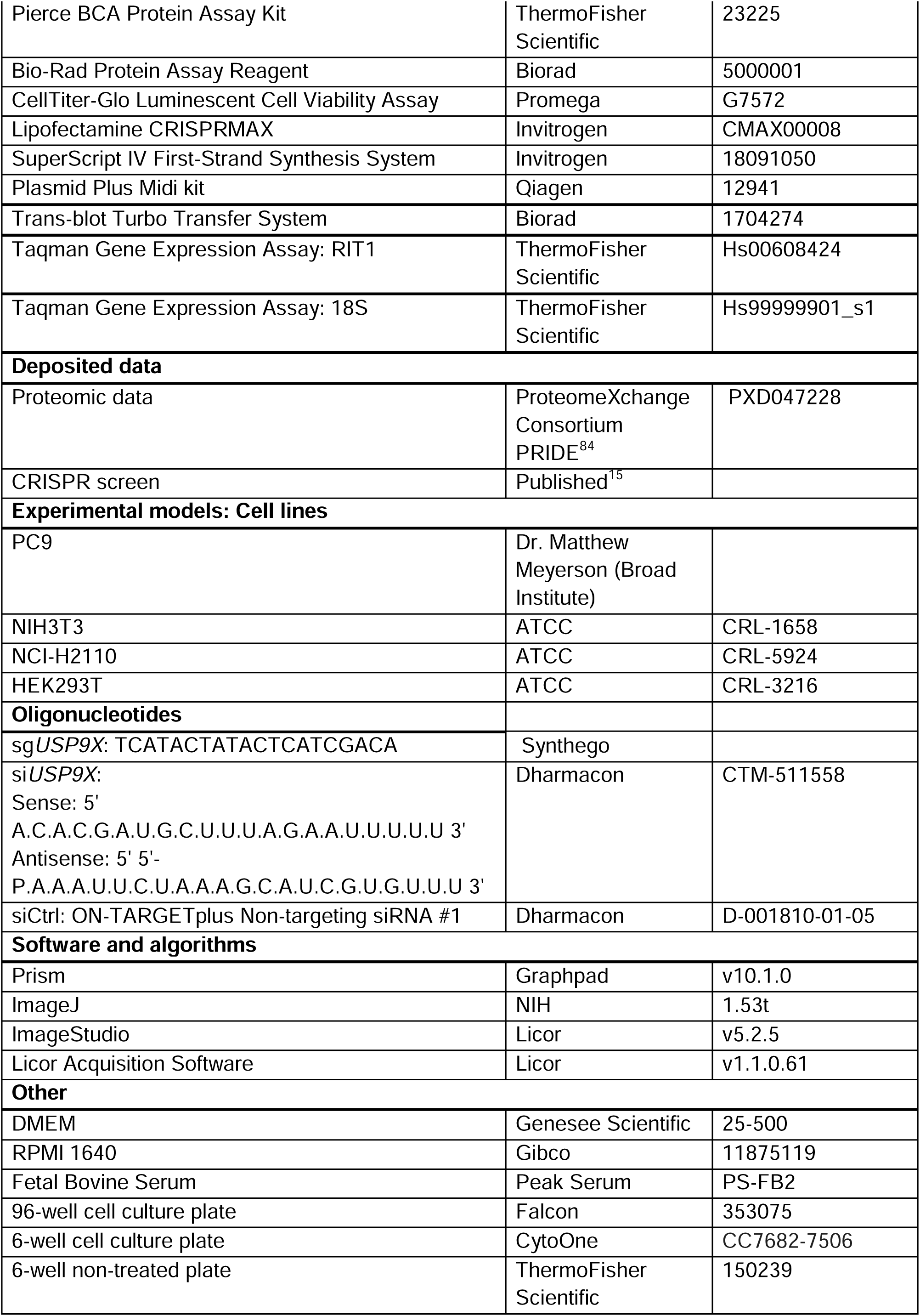
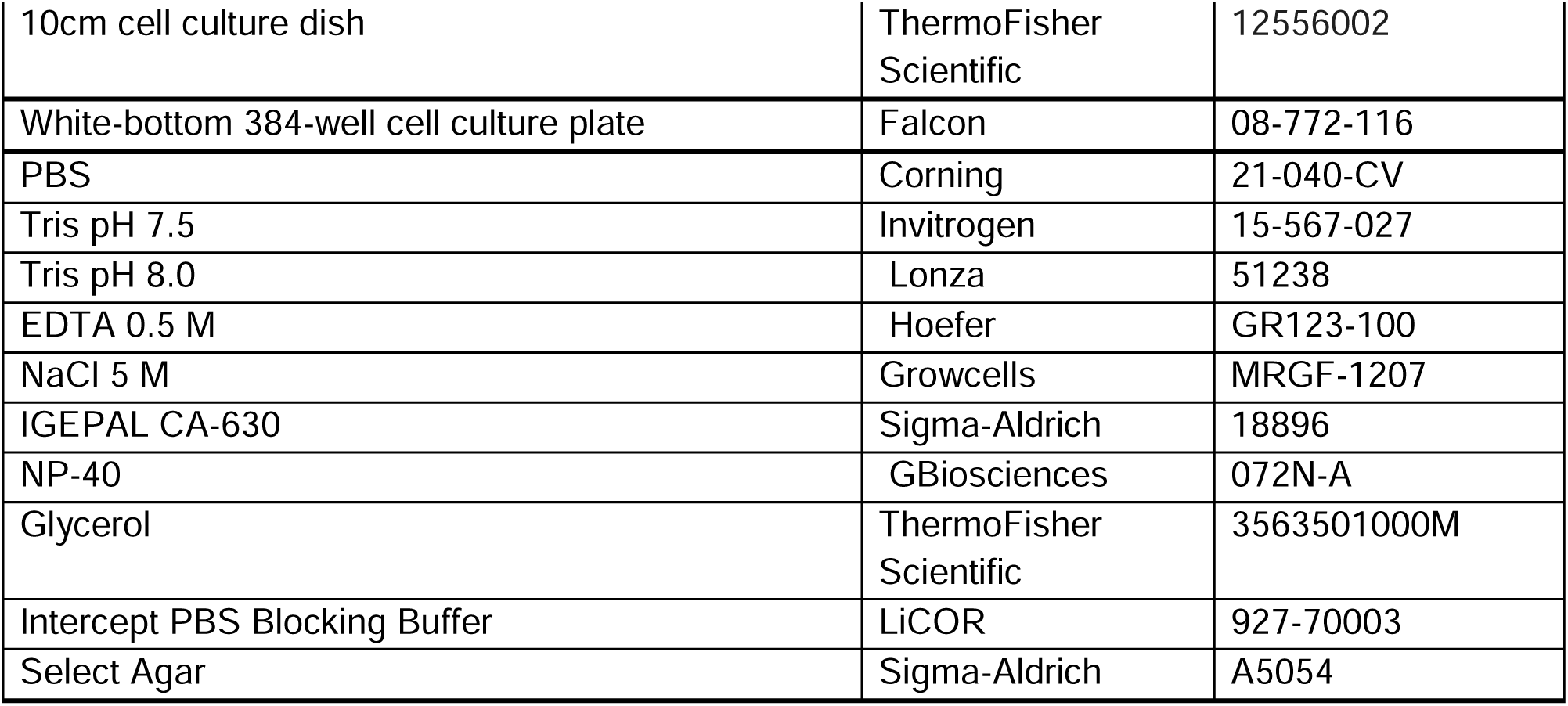

## RESOURCE AVAILABILITY

### Lead contact

Further information and requests for resources and reagents should be directed to and will be fulfilled by the lead contact, Dr. Alice Berger.

### Materials availability

This study did not generate new unique reagents. Reagents used in this study are commercially available or available upon request to the lead author.

### Data and code availability

● The data supporting the findings of this study are available within the article and its supplementary materials.
● The mass spectrometry proteomics data have been deposited to the ProteomeXchange Consortium via the PRIDE^84^ partner repository with the dataset identifier PXD047228.
● Analysis of CRISPR screen data is based on a previously published manuscript^15^.
● This study did not generate code. The custom macro for image analysis is available upon request.
● Any additional information required to reanalyze the data reported in this paper is available from the lead contact (Dr. Alice Berger) upon request.

## EXPERIMENTAL MODEL AND STUDY PARTICIPANT DETAILS

### Cell lines

PC9 cells were a gift from Dr. Matthew Meyerson (Broad Institute). PC9-Cas9 cells were generated as previously described^15^. NIH3T3, NCI-H2110, and HEK293T cells were obtained from ATCC (CRL-1658, CRL-5924, and CRL-3216, respectively). PC9 and NCI-H2110 cells were cultured in RPMI-1640 (Gibco) supplemented with 10% Fetal Bovine Serum (FBS). NIH3T3 and HEK293T cells were cultured in Dulbecco’s Modified Eagle’s Medium (DMEM, Genesee Scientific) supplemented with 10% FBS (Peak Serum, PS-FB2). All cells were maintained at 37°C in 5% CO_2_ and confirmed mycoplasma-free.

## METHOD DETAILS

### Cell line generation

RIT1^M90I^, RIT1^M90I^ + sg*USP9X*, and RIT1^M90I^ + sg*RIT1* cells were generated as previously described^15^. PC9-Cas9 Parental + sg*USP9X* cells were generated by co-transfecting with lipofectamine CRISPR max (Life Technologies) and a synthetic guide RNA: TCATACTATACTCATCGACA

Single cells were plated in a 96-well cell culture plate (Falcon), and clones were expanded and validated by Western blot analysis and Sanger sequencing. H2110iCas9 cells were generated as previously described^15^.

### Transformation and plasmid preparation

To propagate plasmids, One Shot TOP10 Chemically Competent *E. coli* (ThermoFisher Scientific) were transformed with 1 μg of plasmid. Bacteria were propagated and plasmid was isolated using the Plasmid Plus Midi kit (Qiagen) as per the manufacturer’s protocol.

### siRNA treatment

Lyophilized siRNAs were ordered from Dharmacon and resuspended to make 100 μM stock solutions. For siCtrl conditions, ON-TARGETplus Non-targeting siRNA #1 was used. The sequence for si*USP9X* is as follows:

Sense: 5’ ACACGAUGCUUUAGAAUUUUU 3’

Antisense: 5’ PAAAUUCUAAAGCAUCGUGUUU 3’

Transfections were performed using Lipofectamine RNAiMAX (Life Technologies) following the manufacturer’s protocol.

### Dose response curves

For drug treatment experiments in PC9 cells, cells were plated in white-bottom 384-well plates (Falcon) at a density of 400 cells per well in 40 μL of media. For siRNA experiments, 1×10^6^ cells were plated in a 10 cm cell culture dish (ThermoFisher Scientific). 24 hours later, cells were transfected with 120 pmol of siCtrl or si*USP9X* (Dharmacon) following the Lipofectamine RNAiMAX transfection procedure (Life Technologies). 48 hours later, cells were plated in 384-well plates as described above. 24 hours after cell plating, a serial dilution of erlotinib or osimertinib was performed using a D300e dispenser (Tecan). 72 hours post-treatment, 10 μL of CellTiterGlo reagent (Promega) was added to each well and luminescence was quantified on an Envision MultiLabel Plate Reader (PerkinElmer). The viable cell fraction was calculated by comparing the viability of drug-treated cells to the average viability of cells treated with DMSO only (Sigma-Aldrich), normalized by fluid volume. Curve fitting was performed using GraphPad Prism (v10.1.0). Inhibitors were obtained from SelleckChem: Erlotinib-OSI-774 (S1023) and Osimertinib-AZD92921 (S7297). Area-under-the curve (AUC) analyses were performed in Prism 10 (v10.0.3).

### Cell lysis and immunoblotting

Whole-cell extracts for immunoblotting were prepared by washing cells with cold PBS (Corning) supplemented with phosphatase inhibitors (ThermoFisher) on ice and then scraping cells in RTK lysis buffer [20 mM Tris (pH 8.0), 2 mM EDTA (pH 8.0), 137 mM NaCl, 1% IGEPAL CA-630, 10% Glycerol, and ddH_2_0] supplemented with phosphatase inhibitors (ThermoFisher Scientific) and protease inhibitors (ThermoFisher Scientific, EDTA-free). Lysates were incubated on ice for 20 min. Following centrifugation (13,000 rpm for 20 min), lysates were quantified using the Pierce BCA Protein Assay Kit (ThermoFisher Scientific) in a 96-well plate and read on an Accuris Smartreader 96 (MR9600). Lysates were separated by SDS-PAGE and transferred to PVDF membranes using the Trans-blot Turbo Transfer System (BioRad). Membranes were blocked in Intercept PBS blocking buffer (LiCOR) for 1 h at room temperature followed by overnight incubation at 4°C with primary antibodies diluted in blocking buffer. IRDye (LiCOR) secondary antibodies were used for detection and were imaged on the LiCOR Odyssey DLx. Images were acquired using the Licor Acquisition Software (v1.1.0.61) from LiCOR Biosciences. Loading control and experimental proteins were probed on the same membrane unless indicated otherwise. For clarity, loading control is shown below experimental conditions in all panels regardless of the relative molecular weights of the experimental protein(s). Quantification and normalization of Western blot band intensity was performed following protocols from Invitrogen (iBright Imaging Systems).

For immunoprecipitation experiments in Figures 4D and 4F-G: cells were lysed in EBC buffer (50 mM Tris pH 7.5, 120 mM NaCl, 0.5% NP-40/IGEPAL CA-630) supplemented with protease inhibitors (Thermo Scientific) and phosphatase inhibitors (Thermo Scientific). To prepare the Whole Cell Lysates (WCL), 3 × SDS sample buffer was directly added to the cell lysates and sonicated before being resolved on SDS-PAGE and subsequently immunoblotted with primary antibodies. The protein concentrations of the lysates were measured using the Bio-Rad protein assay reagent on a Bio-Rad Model 680 Microplate Reader. For immunoprecipitation, 1 mg lysates were incubated with the appropriate agarose-conjugated primary antibody for 3-4 h at 4°C or with unconjugated antibody (1-2 mg) overnight at 4°C followed by 1 h incubation with Protein G Sepharose beads (GE Healthcare). Immuno-complexes were washed four times with NETN buffer (20 mM Tris, pH 8.0, 100 mM NaCl, 1 mM EDTA and 0.5% NP-40) before being resolved by SDS-PAGE and immunoblotted with indicated antibodies.

Primary antibodies used for immunoblotting: β-Actin 1:1000 (Cell Signaling Technology, 4970), USP9X 1:500 (Proteintech, 55054-1-AP), RIT1 1:1000 (Abcam, Ab53720), Vinculin 1:1500 (Sigma-Aldrich, V9264), Cyclophilin A 1:1000 (Bio-Rad, VMA00535), Cas9 1:1000 (Cell Signaling Technology, 14697), CDC20 1:2000 (Santa Cruz, 13162), Tubulin 1:2000 (Sigma-Aldrich, T5168), Flag 1:2000 (Sigma-Aldrich, F1804), HA 1:2000 (Biolegend, 901503).

### Proliferation assay

PC9-Cas9 cells (Parental, RIT1^M90I^, RIT1^M90I^ + sg*USP9X*, and RIT1^M90I^ + sg*RIT1*) were seeded in triplicate in 6-well tissue culture-treated dishes (CytoOne) at a density of 1×10^5^ cells per well. Cells were counted and passaged every 2-4 days and replated at a density of 1×10^5^ cells per well. Cumulative population doublings were calculated in excel, and statistical analyses were performed in Prism (v10.1.0).

### Soft agar assays

For soft agar colony formation assays, 4×10^5^ PC9-Cas9 cells (Parental, RIT1^M90I^, RIT1^M90I^ + sg*USP9X*, and RIT1^M90I^ + sg*RIT1*) were suspended in 1.3 mL media and 2.7 mL 0.5% select agar (Sigma-Aldrich) in RPMI+10% FBS. 1 mL of this cell suspension was plated into 3 wells on a bottom layer of 0.5% select agar in RPMI+10% FBS in 6-well non-tissue culture treated dishes (Thermo Scientific). For soft agar inhibitor experiments, erlotinib was suspended in the top agar solution at a final concentration of 500 nM. DMSO control conditions were prepared to normalize by DMSO volume. After 10 days of growth, brightfield images were acquired on an ImageExpress (Molecular Devices) microscope using a 4x/0.2 NA objective. Fields of view were tiled in a 9×9 grid to cover the entire well with no overlap. A z-stack with 1125 μm range and 25 μm step size were acquired for each field of view and saved as a 2D minimum projection. All images were analyzed in ImageJ (1.53t) using a custom macro (available upon request). Images were excluded if they were obstructed by the 6-well plate and/or if the agar contained bubbles or other abnormalities.

### Cycloheximide-chase

PC9-Cas9 Parental and sg*USP9X* cells treated with 100 μg/mL cycloheximide (CHX) for the indicated time periods before harvesting protein for Western blot. Cycloheximide (Tocris Bioscience, Cat. No. 0970, CAS No. 66-81-9), was dissolved in DMSO as 100 mg/ml stock solution freshly before each use.

### RT-qPCR

Total RNA was extracted from two biological replicates of parental H2110iCas9 cells and H2110iCas9 + sg*USP9X* cells treated with siCtrl or si*USP9X*. Reverse transcription was performed with 1 μg RNA and the SuperScript IV First-Strand Synthesis System (Invitrogen). 20 ng of cDNA was used for each RT-PCR reaction. cDNA was amplified using TaqMan gene expression assays (ThermoFisher Scientific): RIT1 (assay Hs00608424_m1) and 18S (assay Hs99999901_s1). Reactions were run on the BioRad CFX384 Real-Time system. Expression was normalized to 18S within each sample in the same experiment, and relative expression was quantified using the 2^-ΔΔCt^ method.

### Co-immunoprecipitation

For RIT1^∼Ub^ pulldown, 2 million HEK293T cells were plated in 10 cm cell culture dishes (ThermoFisher Scientific). The next day, cells were transfected with 10 μg of indicated plasmids and jetPRIME reagent (Polyplus), following the manufacturer’s protocol. 24 hours post-transfection, HA pulldown was performed using EZview Red Anti-HA Affinity Gel (Millipore Sigma). In brief, cells were washed with cold PBS supplemented with phosphatase inhibitors (ThermoFisher Scientific). Cells were scraped in 1 mL NP40 lysis buffer (150 mM NaCl, 1% NP40, 10% glycerol, 10mM Tris (pH 8.0), and ddH_2_0) supplemented with protease and phosphatase inhibitors (ThermoFisher Scientific). 50 μL of lysate was reserved for the Whole Cell Lysate. 30 μL of pre-washed EZview beads were added per condition, and samples were incubated for 2 hours at 4°C with shaking. Samples were washed 3X and then prepared for SDS-PAGE and Western blot as described above. For RIT1^∼Ub^ pull downs used for affinity purification mass spectrometry, a similar protocol was used with few substitutions. First, the number of cells was scaled up to 15 million in two 15 cm plates and magnetic anti-HA beads (ThermoFisher Scientific) were used instead. After washing beads with lysis buffer, two additional washes were performed using PBS to remove residual detergent present in the beads. For each experimental condition, four biological replicates were used.

For IP:RIT1 experiments with endogenous USP9X, 3 million HEK393T cells were plated in 10 cm cell culture dishes (ThermoFisher Scientific). The next day, cells were transfected with 2.5 μg of GST-tagged RIT1 plasmids or GFP control plasmid using Lipofectamine 3000 Reagent (ThermoFisher Scientific) following the manufacturer’s protocol. 24 hours later, cells were washed with cold PBS supplemented with phosphatase inhibitors (ThermoFisher Scientific). Cells were scraped in 700 μL lysis buffer (50mM Tris (pH 7.5), 1% IGEPAL-CA-630) supplemented with protease and phosphatase inhibitors (ThermoFisher Scientific). Protein lysates were quantified using the Pierce BCA Protein Assay Kit (ThermoFisher Scientific), and 1 mg of protein was used for each IP condition. 50 μg of protein was set aside for the Whole Cell Lysate. For IP conditions, each lysate was pre-cleared with 20 μL of pre-washed Protein A agarose beads (Cell Signaling 9863S). Next, 1 μg of RIT1 antibody (Abcam Ab53720) or 1 μg of control rabbit IgG antibody (R&D Systems AB-105-C) was added, and samples were incubated for 2 hours at 4°C with shaking. 20 μL of beads were added to each tube, and samples were incubated overnight at 4°C with shaking. The next day, samples were washed 3X and prepared for SDS-PAGE and Western blot as described above.

### *In vitro* ubiquitination assay

HEK293T cells were transfected with RIT1, USP9X, and His-ubiquitin constructs. 36 hours after transfection, 10 μM MG132 was added to block proteasome degradation, and cells were harvested in denatured buffer (6 M guanidine-HCl, 0.1 M Na_2_HPO_4_/NaH_2_PO_4_, 10 mM imidazole), followed by Ni-NTA (Ni-nitrilotriacetic acid) purification and immunoblot analysis. MG-132 (Selleck Chemicals, Cat. No. S2619, CAS No. 1211877-36-9), was dissolved in DMSO as a 10 mM stock solution and stored in -20°C.

### Affinity purification/mass spectrometry

On-bead trypsin digests were performed as previously described^14^, and digested tryptic peptides were analyzed by LC-MS/MS on Orbitrap Fusion Lumos Tribrid Mass Spectrometer (Thermo Fisher Scientific) using the same configuration and settings as previously reported^85^. Acquired MS data were analyzed using a workflow previously described^85,86^. Briefly, spectra were searched in Protein Prospector (version 6.2.4^87^) against human proteome (SwissProt database downloaded on 01/18/2021) and decoy database of corresponding randomly shuffled peptides. Search engine parameters were as follows: “ESI-Q-high-res” for the instrument, trypsin as the protease, up to 2 missed cleavages allowed, Carbamidomethyl-C as constant modification, default variable modifications, up to 3 modifications per peptide allowed, 15 ppm precursor mass tolerance, and 25 ppm tolerance for fragment ions. The false discovery rate was set to <1% for peptides, and at least 3 unique peptides per protein were required. Protein Prospector search results formatted as BiblioSpec spectral library were imported into Skyline (v21) to quantify peptide and protein abundances using MS1 extracted chromatograms^88^. Statistical analysis of observed protein abundances was performed using MSstats package integrated in Skyline^89^.

Abundance per peptide represents the log_2_-abundance of the peak intensity (AUC) from mass-spectrometry for each peptide. Mean per-peptide abundance is the average enrichment of abundance for each peptide across replicates. Mean of mean-peptide abundance was calculated as the average of mean-peptide abundance for all peptides for the protein across repeats. Mean of mean-peptide abundance was used to generate the heatmap (**Figure S4A**). Heatmap shows a subset of those proteins with at least a 5-fold enrichment over empty vector (EV).

### CRISPR data analysis

CRISPR scores were calculated as previously described^15^. In brief, all data were scaled so that the median of non-essential genes (based on previously published lists in DepMap^90^) is 0 and the median of essential genes is -1. CRISPR scores were defined as this scaled, normalized log-fold-change data. All data from this CRISPR screen are available within the main figures and supplemental information from the associated published manuscript^15^.

### DepMap analyses

For correlation analyses, data were explored in the Cancer Dependency Map (DepMap) online portal (https://depmap.org/portal/). Proteomics data were captured from^36^ and available within DepMap. RIT1, CDC20, and USP9X expression were evaluated in the Expression Public 23Q2 datasets and previously published^37^. Correlation and statistical tests were performed in Prism (v10.1.0).

## QUANTIFICATION AND STATISTICAL ANALYSIS

Data are expressed as mean ± s.d. unless otherwise noted. Exact numbers of biological and technical replicates for each experiment are reported in the Figure Legends. p-values less than 0.05 were considered statistically significant based on the appropriate statistical test for the experiment in question. For all data, *p<0.05, **p<0.01, ***p<0.001, ****p<0.0001. Data were analyzed using Prism Software 10.0 (GraphPad).

