## Supplemental Figures and Legends for "The deubiquitinase USP9X regulates RIT1 protein abundance and oncogenic phenotypes"

**SUPPLEMENTAL INFORMATION**

**Figure S1.**

**A,** Box plots showing average CRISPR score of indicated sgRNAs in PC9-Cas9 Parental (+Luciferase/Control) cells or $RIT1^{M90I}$-mutant PC9-Cas9 cells. Box plots show the median (center line) and the min and max range of replicates. For Control conditions, n = 3 biological replicates. For $RIT1^{M90I}$-mutant cells, n = 2 biological replicates. p-values calculated by unpaired two-tailed t-tests. **B,** Dose-response curves of PC9-Cas9 Parental cells and $RIT1^{M90I}$-mutant PC9-Cas9 cells with indicated gene knockouts (sg*RIT1* and sg*USP9X*) treated with osimertinib for 72 hours. CellTiterGlo was used to quantify viable cell fraction determined by normalization to DMSO control. Data shown are the mean $\pm$ s.d. of two technical replicates. Data are representative results from n = 2 independent experiments. **C,** Area-under-the-curve (AUC) analysis of dose response curves shown in (D). p-values calculated by unpaired two-tailed t-tests. **D,** Dose-response curve of $RIT1^{M90I}$-mutant PC9 cells treated with siCtrl or si*USP9X* for 48 hours, prior to treatment with osimertinib for 72 hours. CellTiterGlo was used to quantify viable cell fraction determined by normalization to DMSO control. Data shown are the mean $\pm$ s.d. of two technical replicates. Data are representative results from n = 3 independent experiments.

**Figure S2.**

Proliferation of PC9-Cas9 Parental cells and $RIT1^{M90I}$-mutant PC9-Cas9 cells with indicated gene knockouts (sg*RIT1* and sg*USP9X*) treated with DMSO (vehicle). Data shown are the mean $\pm$ s.d. of three technical replicates per cell line. Data are representative results from n = 2 independent experiments. p-value calculated by multiple unpaired two-tailed t-tests.

**Figure S3.**

**A,** Western blot of NCI-H2110 cells treated with indicated siRNAs for 72 hours. Vinculin serves as a loading control. **B,** Quantification of Western blot bands based on (A) and additional replicates. Data shown are the mean $\pm$ s.d. of three independent experiments with 2-3 technical replicates per condition. p-value calculated by paired two-tailed t-test. **C,** Western blot of NCI-H2110iCas9 cells treated with 1 µg/mL Dox for 7 days to induce Cas9 expression. Cyclophilin A serves as a loading control. **D,** Quantification of Western blot bands in (C). Data shown are the mean $\pm$ s.d. of two independent experiments. p-value calculated by paired two-tailed t-test. **E,** Relative expression of *RIT1* as determined by qPCR and ΔΔCt analysis. Data shown are the mean $\pm$ s.d. of three technical replicates per condition. Data are representative of results from n = 2 independent experiments. ns = not significant by paired two-tailed t-test.

**Figure S4.**

**A,** Heatmap of peptides detected in affinity purification/mass spectrometry (AP/MS) experiment. Data were filtered for proteins that were at least 5 times higher in $RIT1^{\sim Ub}$ condition compared to EV (Empty Vector). Abundance values are the $log_2$-based number of the peak intensity from the MS. The mean of all peptides was combined across biological replicates. For EV samples, n = 7 biological replicates. For $RIT1^{\sim Ub}$ samples, n = 4 biological replicates. **B,** Abundance ($log_2$-transformed) of individual RIT1 peptides in Empty Vector (EV) control and $RIT1^{\sim Ub}$ conditions from AP/MS. **C,** Abundance ($log_2$-transformed) of individual LZTR1 peptides in EV and $RIT1^{\sim Ub}$ conditions from AP/MS. **D,** Abundance ($log_2$-transformed) of individual USP9X peptides in EV and $RIT1^{\sim Ub}$ conditions from AP/MS. For B-D, some peptides were detected in all replicates while other peptides were only detected in some replicates.

**Supplementary Table 1. Deubiquitinases and E3 ligases identified in whole-genome screen of *RIT1*-mutant PC9 cells.**

ΔCRISPR Score of all genes listed in gene rank plot in Figure 1B. DUBs are highlighted in red and E3 ligases are highlighted in green.

Figure S1.

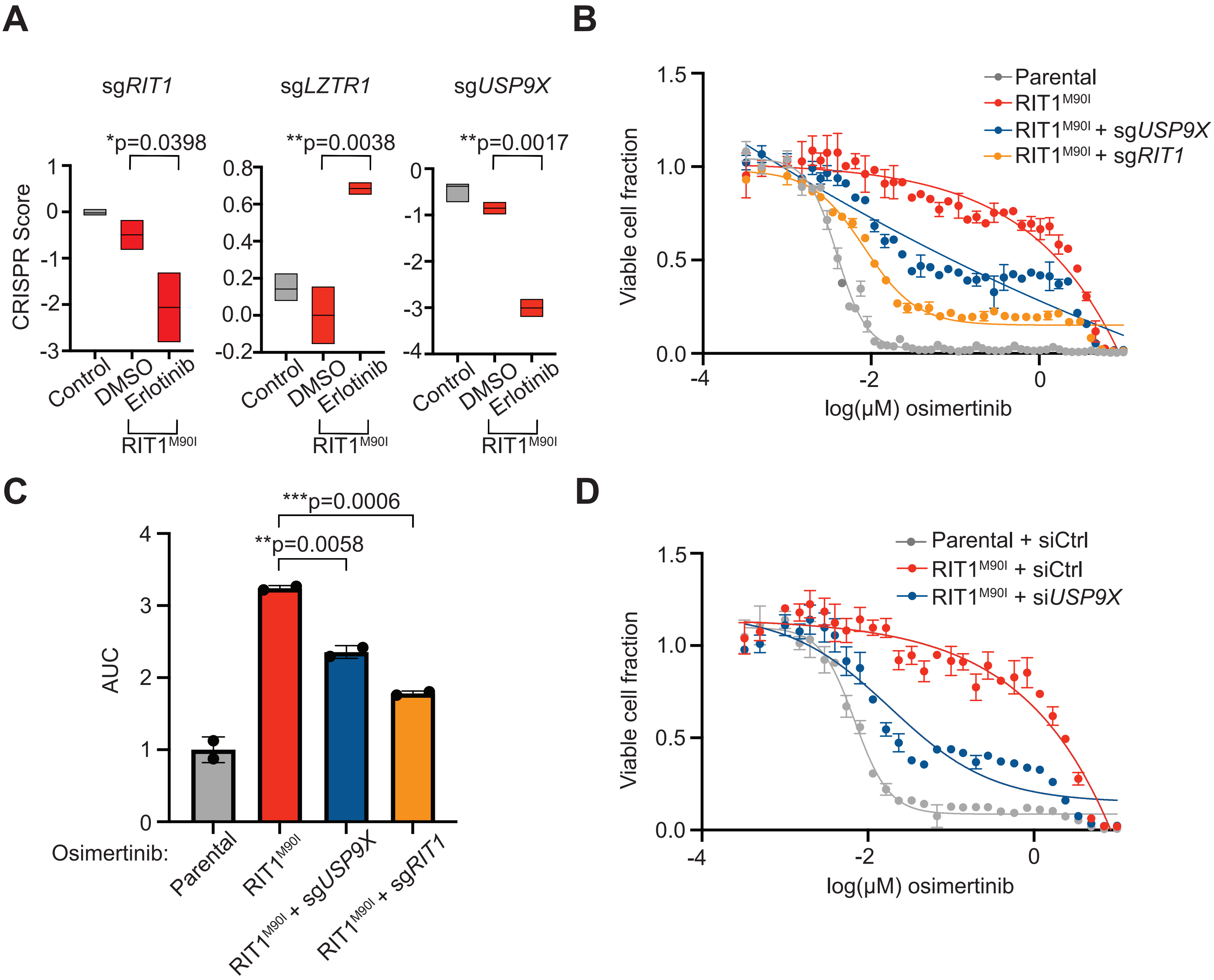

A
sgRIT1
sgLZTR1
sgUSP9X
*p=0.0398
**p=0.0038
**p=0.0017
CRISPR Score
Control
DMSO
Erlotinib
RIT1M90I
B
Parental
RIT1M90I
RIT1M90I + sgUSP9X
RIT1M90I + sgRIT1
Viable cell fraction
log(μM) osimertinib
C
***p=0.0006
**p=0.0058
AUC
Osimertinib:
Parental
RIT1M90I
RIT1M90I + sgUSP9X
RIT1M90I + sgRIT1
D
Parental + siCtrl
RIT1M90I + siCtrl
RIT1M90I + siUSP9X
Viable cell fraction
log(μM) osimertinib

**Figure S2.**

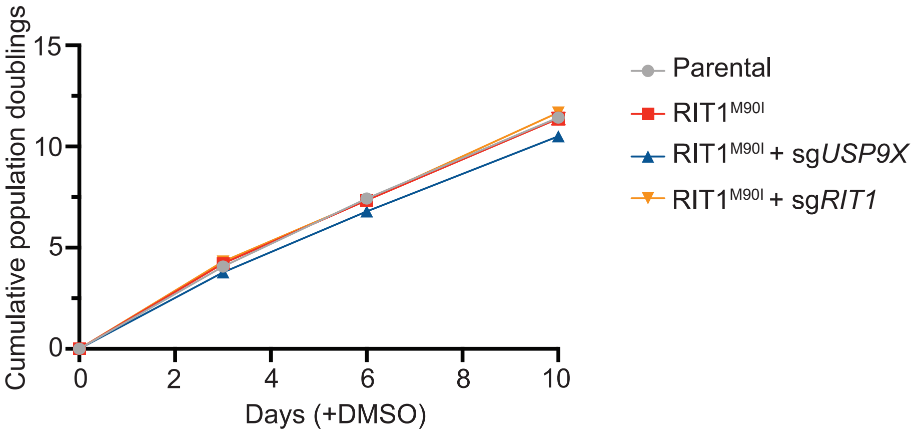
Cumulative population doublings
15
10
5
0
0
2
4
6
8
10
Days (+DMSO)
Parental
RIT1$^{M90I}$
RIT1$^{M90I}$ + sg*USP9X*
RIT1$^{M90I}$ + sg*RIT1*

**Figure S3.**

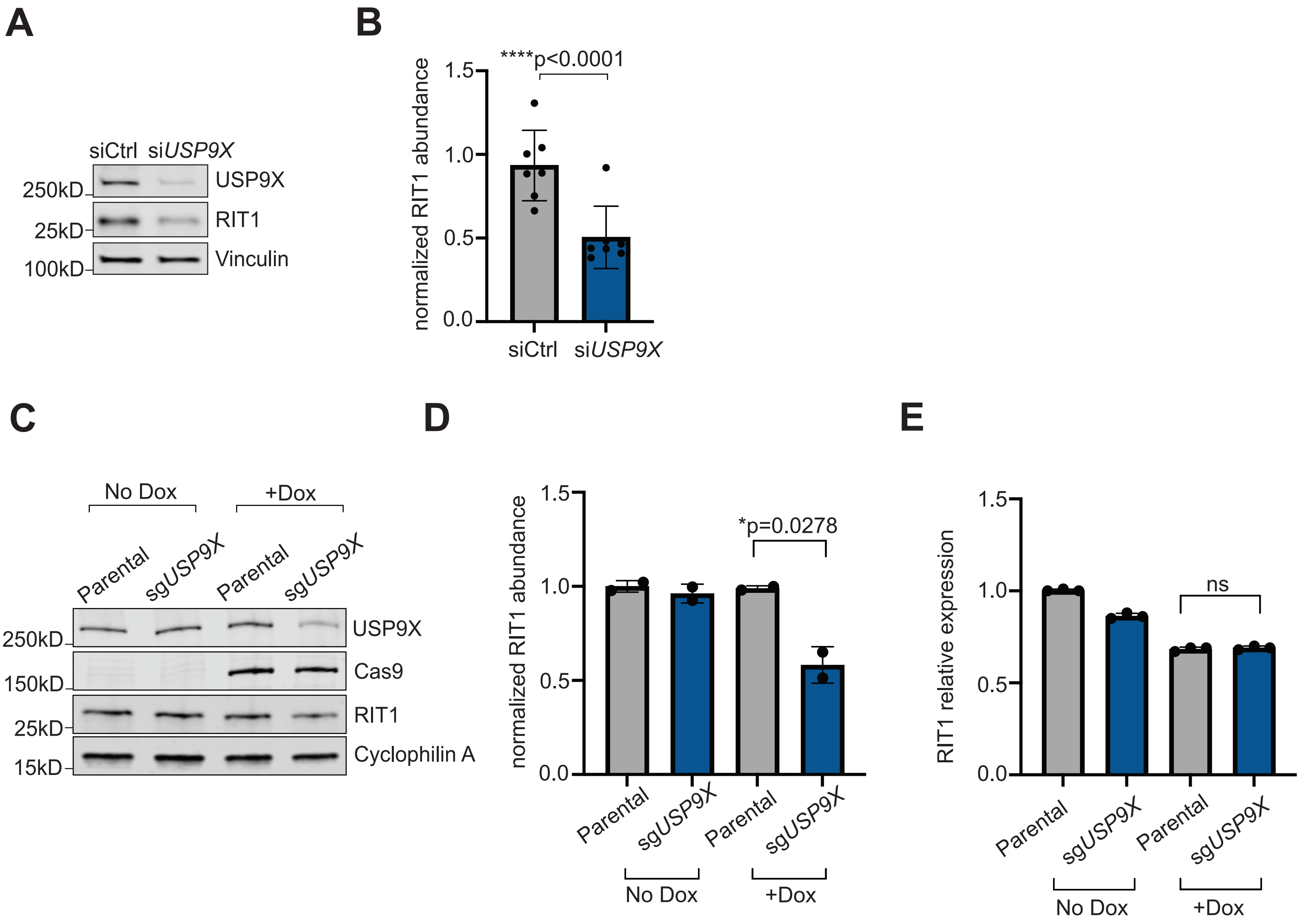
A
siCtrl
siUSP9X
250kD
USP9X
25kD
RIT1
100kD
Vinculin
B
****p<0.0001
normalized RIT1 abundance
1.5
1.0
0.5
0.0
siCtrl
siUSP9X
C
No Dox
+Dox
Parental
sgUSP9X
Parental
sgUSP9X
250kD
USP9X
150kD
Cas9
25kD
RIT1
15kD
Cyclophilin A
D
normalized RIT1 abundance
*p=0.0278
1.5
1.0
0.5
0.0
Parental
sgUSP9X
Parental
sgUSP9X
No Dox
+Dox
E
RIT1 relative expression
ns
1.5
1.0
0.5
0.0
Parental
sgUSP9X
Parental
sgUSP9X
No Dox
+Dox

**Figure S4.**

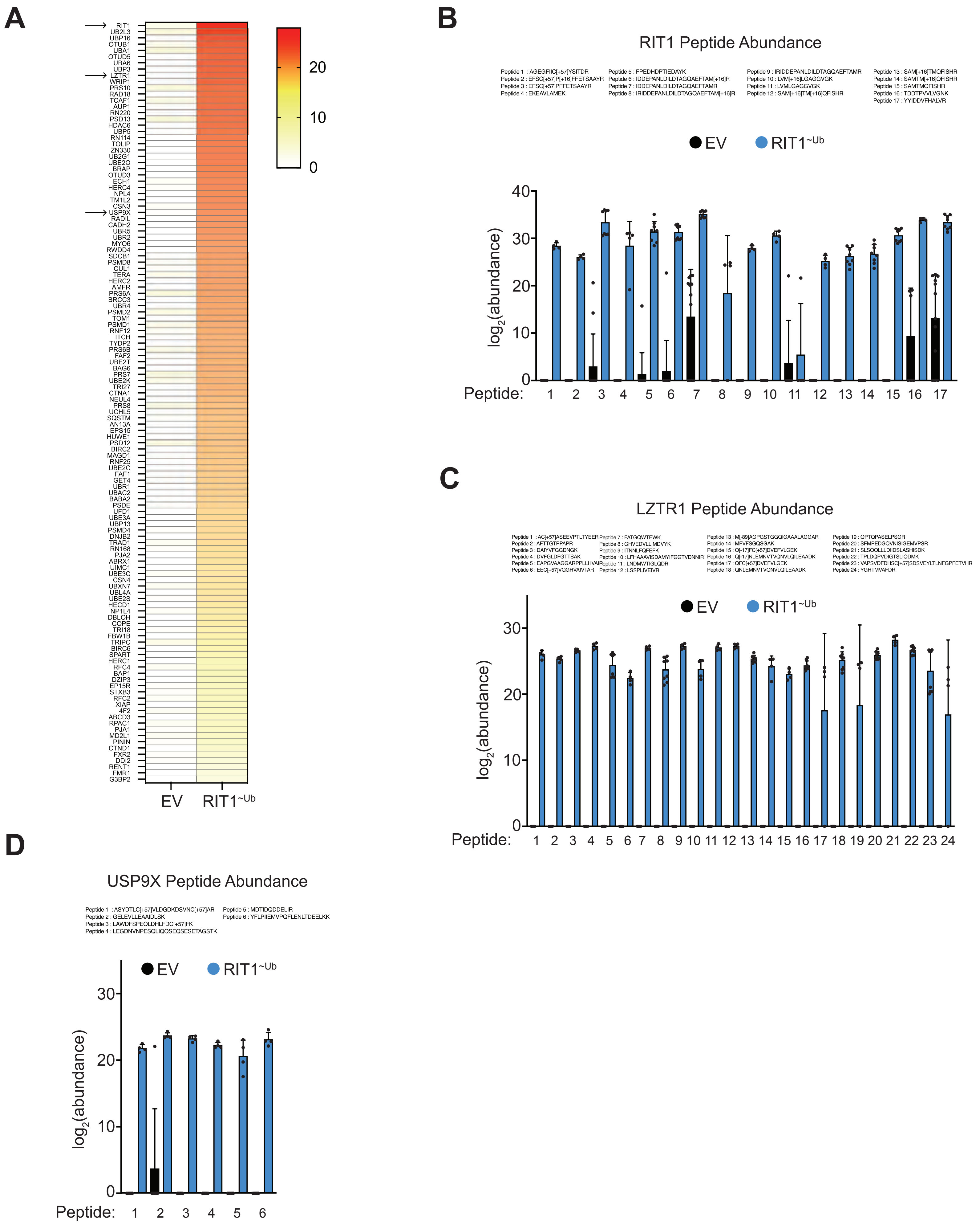
A
RIT1
UB2L3
UBP16
OTUB1
UBA1
OTUD5
UBA6
UBP3
LZTR1
WRIP1
PRS10
RAD18
TCAF1
AUP1
RN220
PSD13
HDAC6
UBP5
RN114
TOLIP
ZN330
UB2G1
UBE2O
BRAP
OTUD3
ECH1
HERC4
NPL4
TM1L2
CSN3
USP9X
RADIL
CADH2
UBR5
UBR2
MYO6
RWDD4
SDCB1
PSMD8
CUL1
TERA
HERC2
AMFR
PRS6A
BRCC3
UBR4
PSMD2
TOM1
PSMD1
RNF12
ITCH
TYDP2
PRS6B
FAF2
UBE2T
BAG6
PRS7
UBE2K
TRI27
CTNA1
NEUL4
PRS8
UCHL5
SQSTM
AN13A
EPS15
HUWE1
PSD12
BIRC2
MAGD1
RNF25
UBE2C
FAF1
GET4
UBR1
UBAC2
BABA2
PSDE
UFD1
UBE3A
UBP13
PSMD4
DNJB2
TRAD1
RN168
PJA2
ABRX1
UIMC1
UBE3C
CSN4
UBXN7
UBL4A
UBE2S
HECD1
NP1L4
DBLOH
COPE
TRI18
FBW1B
TRIPC
BIRC6
SPART
HERC1
RFC4
BAP1
DZIP3
EP15R
STXB3
RFC2
XIAP
4F2
ABCD3
RPAC1
PJA1
MD2L1
PININ
CTND1
FXR2
DDI2
RENT1
FMR1
G3BP2
20
10
0
EV
RIT1~Ub
B
RIT1 Peptide Abundance
Peptide 1 : AGEGFIIC[+57]YSITDR
Peptide 2 : EFSC[+57]P[+16]FFETSAAYR
Peptide 3 : EFSC[+57]PFFETSAAYR
Peptide 4 : EKEAVLAMEK
Peptide 5 : FPEDHDPTIEDAYK
Peptide 6 : IDDEPANLDILDTAGQAEFTAM[+16]R
Peptide 7 : IDDEPANLDILDTAGQAEFTAMR
Peptide 8 : IRIDDEPANLDILDTAGQAEFTAM[+16]R
Peptide 9 : IRIDDEPANLDILDTAGQAEFTAMR
Peptide 10 : LVM[+16]LGAGGVGK
Peptide 11 : LVMLGAGGVGK
Peptide 12 : SAM[+16]TM[+16]QFISHR
Peptide 13 : SAM[+16]TMQFISHR
Peptide 14 : SAMTM[+16]QFISHR
Peptide 15 : SAMTMQFISHR
Peptide 16 : TDDTPVVLVGNK
Peptide 17 : YYIDDVFHALVR
EV
RIT1~Ub
log2(abundance)
40
30
20
10
0
Peptide: 1 2 3 4 5 6 7 8 9 10 11 12 13 14 15 16 17
C
LZTR1 Peptide Abundance
Peptide 1 : AC[+57]ASEEVPTLTYEER
Peptide 2 : AFTTGTPPAPR
Peptide 3 : DAIYVFGGDNGK
Peptide 4 : DVFGLDFGTTSAK
Peptide 5 : EAPGVAAGGARPPLLHVAIR
Peptide 6 : EEC[+57]VQGHVAIVTAR
Peptide 7 : FATGQWTEWK
Peptide 8 : GHVEDVLLIMDVYK
Peptide 9 : ITNNLFQFEFK
Peptide 10 : LFHAAAVISDAMYIFGGTVDNNIR
Peptide 11 : LNDMWTIGLQDR
Peptide 12 : LSSPLIVEIVR
Peptide 13 : M[-89]AGPGSTGGQIGAAALAGGAR
Peptide 14 : MFVFSGQSGAK
Peptide 15 : Q[-17]FC[+57]DVEFVLGEK
Peptide 16 : Q[-17]NLEMNVTVQNVLQILEAADK
Peptide 17 : QFC[+57]DVEFVLGEK
Peptide 18 : QNLEMNVTVQNVLQILEAADK
Peptide 19 : QPTQPASELPSGR
Peptide 20 : SFMPEDGQVNISIGEMVPSR
Peptide 21 : SLSQQLLLDIIDSLASHISDK
Peptide 22 : TPLDQPVDIGTSLIQDMK
Peptide 23 : VAPSVDFDHSC[+57]SDSVEYLTLNFGPFETVHR
Peptide 24 : YGHTMVAFDR
EV
RIT1~Ub
log2(abundance)
30
20
10
0
Peptide: 1 2 3 4 5 6 7 8 9 10 11 12 13 14 15 16 17 18 19 20 21 22 23 24
D
USP9X Peptide Abundance
Peptide 1 : ASYDTLC[+57]VLDGDKDSVNC[+57]AR
Peptide 2 : GELEVLLEAAIDLSK
Peptide 3 : LAWDFSPEQLDHLFDC[+57]FK
Peptide 4 : LEGDNVNPESQLIQQSEQSESETAGSTK
Peptide 5 : MDTIDQDDELIR
Peptide 6 : YFLPIIEMVPQFLENLTDEELKK
EV
RIT1~Ub
log2(abundance)
30
20
10
0
Peptide: 1 2 3 4 5 6
